# Sampling bias exaggerates a textbook example of a trophic cascade

**DOI:** 10.1101/2020.05.05.079459

**Authors:** Elaine M. Brice, Eric J. Larsen, Daniel R. MacNulty

## Abstract

Understanding how ecosystems respond to the loss and recovery of large predators is a major challenge because these free-living systems are difficult to sample properly. We show how an accepted practice of nonrandom sampling has confounded this understanding in a textbook system (Yellowstone National Park) where carnivore [*Canis lupus* (wolf)] recovery is often associated with a trophic cascade involving changes in herbivore [*Cervus canadensis* (elk)] behavior and density that promote plant regeneration. Long-term data indicate that a customary practice of sampling only the tallest young plants overestimated regeneration of overstory aspen (*Populus tremuloides*) by a factor of 3-8 compared to random sampling. Sampling only the tallest young plants favored plants taller than the preferred browsing height of elk and overlooked non-regenerating aspen stands. Our results demonstrate how seemingly minor departures from principled sampling can generate substantial misunderstandings about the strength of trophic cascades in response to large predator recovery.

## INTRODUCTION

Knowledge about the occurrence and strength of trophic cascades (indirect effects of predators on plants and abiotic processes) is vital to understand the forces that structure food webs and to guide the practices of ecosystem conservation, restoration, and rewilding. Much of the empirical information about trophic cascades derives from tractable systems that are variously small scale, invertebrate, and captive (Ford & Goheen 2015; Piovia-Scott *et al*. 2017; Alston *et al*. 2019). Substantially less is known about trophic cascades in free-living large vertebrate systems due in part to the difficulty and cost of measuring such systems in accordance with basic sampling principles (Ford & Goheen 2015; Allen *et al*. 2017; Hayward *et al*. 2019).

Concern about sampling bias is a core issue in the longstanding debate about the cascading effects of reintroduced wolves in northern Yellowstone National Park (YNP; Bilyeu *et al*. 2008; Kauffman *et al*. 2013; Peterson *et al*. 2014; Winnie 2014; Fleming 2019; Hayward *et al*. 2019). In this system, the trophic cascade hypothesis is that wolves (*Canis lupus*) scared away and/or killed enough elk (*Cervus canadensis*) to allow woody deciduous plants (*Populus* spp., *Salix* spp.) to recover from decades of unchecked browsing. A primary support for this hypothesis is time series data showing annual decreases in browsing and annual increases in plant height following wolf reintroduction (Beschta & Ripple 2016; Beschta *et al*. 2018; Painter *et al*. 2018; Painter & Tercek 2020). A negative correlation between browsing and plant height is considered critical evidence of a trophic cascade because it demonstrates the mechanism connecting the lower two trophic levels: reduced browsing increases plant growth, leading to escape from browsing if plants grow tall (Beyer *et al*. 2007; Beschta & Ripple 2016; Beschta *et al*. 2018). The traditional, though untested, assumptions are that (*i*) plants taller than 200 cm escape the reach of elk and recruit into the aspen overstory, and (*ii*) plants shorter than 200 cm are browsed with equal intensity.

Studies of comparable systems suggest that plant height has a nonlinear effect on elk browsing, such that browsing peaks at a preferred height of < 200 cm (Rounds 1979; Motta 2003; Renaud *et al*. 2003; Konôpka *et al*. 2018; Maxwell *et al*. 2019). If plant height beyond this preference reduces browsing, a negative browsing-plant height correlation may not reflect indicate a trophic cascade (Peterson *et al*. 2014). The more typical concern is that reports of decreased browsing and increased plant height after wolf reintroduction are based on biased sampling (Kauffman *et al*. 2013; Peterson *et al*. 2014; Winnie 2014; Fleming 2019).

Nonrandom sampling underpins nearly every published annual trend in browsing and height of young aspen (*Populus tremuloides*) that has been attributed to a cascading effect of wolves (Table 1). In this case, nonrandom sampling is the practice of measuring the three or five tallest young aspen within a stand. Five tallest data comprise most trends and they originate from one of two time series (Ripple & Beschta 2007; Painter *et al*. 2014). A third time series uses three tallest data (Halofsky *et al*. 2008). All three time series were built from a single year of sampling by retrospectively inferring the past browsing and height of sampled aspen using plant architecture methods (Keigley & Frisina 1998). Only one time series was built by annually sampling randomly selected young aspen: an unpublished dataset from E. Larsen that has received limited attention (Peterson *et al*. 2014, 2020).

**Table 1.** Peer-reviewed publications showing annual trends in height and (or) browsing of young aspen in Yellowstone National Park linked to the cascading effects of wolves. Listed are the authors and publication year (Authors [Year]), number of the relevant data figure in the article (Fig.), source of data shown in relevant data figure (Data Source), timespan covered by the data (Data Years), type of data collected (Height, Herbivory), and method of data collection (selective sampling of the tallest young aspen [Tallest] or random sampling of all young aspen [Random]). Checkmarks indicate which data were collected and with which sampling method. Shaded cells indicate articles that reproduced or extended data originating in Ripple and Beschta (2007), and dashed-outlined cells indicate articles that reproduced or extended data originating in Painter et al. (2014). Underlined data in Peterson et al. (2014, 2020) were unpublished data from E. Larsen, and are the subject of this article.

| Authors (Year) | Fig. | Data Source | Data Years | Data Type |  | Sampling Method |  |
| --- | --- | --- | --- | --- | --- | --- | --- |
|  |  |  |  | Height | Browsing | Tallest* | Random |
| Ripple & Beschta (2007) | 1c-d | Original data (98 stands) | 1998-2006 | ✓ | ✓ | ✓ |  |
| Beschta & Ripple (2009) | 8b | Ripple & Beschta (2007) | 1999-2005 | ✓ | ✓ | ✓ |  |
| Ripple & Beschta (2012) | 1c-d | Ripple & Beschta (2007)<br>Original data (resampled 98 stands) | 1998-2006<br>2010 | ✓ | ✓ | ✓ |  |
| Beschta & Ripple (2016) | 3g | Ripple & Beschta (2007, 2012) | 1998-2006, 2010 | ✓ |  | ✓ |  |
| Beschta et al. (2018) | 2b-c | Ripple & Beschta (2007, 2012)<br>Original data (60 additional stands) | 1999-2006, 2010<br>2005-2015 | ✓ | ✓ | ✓ |  |

Table 1 continued
| Authors (Year) | Fig. | Data Source | Data Years | Data Type |  | Sampling Method |  |
| --- | --- | --- | --- | --- | --- | --- | --- |
|  |  |  |  | Height | Browsing | Tallest* | Random |
| Painter et al. (2014)<sup>†</sup> | 7a | Original data (87 stands) | 2003-2012 | ✓ | ✓ | ✓ |  |
| Peterson et al. (2014) | 1c | Painter et al. (2014)<br><u>Original data (113 stands)</u> | 2003-2012<br>1999-2013<sup>‡</sup> | ✓ |  | ✓ | ✓ |
| Painter et al. (2015)<sup>†</sup> | 2a | Painter et al. (2014) | 2003-2012 | ✓ |  | ✓ |  |
| Beschta et al. (2016) | 5c | Painter et al. (2014) | 2002<sup>§</sup>-2012 |  | ✓ | ✓ |  |
| Painter et al. (2018)<sup>†</sup> | 3a | Ripple & Bestchta (2007, 2012)<br>Painter et al. (2014) | 1999-2006, 2010<br>2005-2012 |  | ✓ | ✓ |  |
| Halofsky et al. (2008) | 3a-b | Original data (44 stands) | 1995-2004 | ✓ | ✓ | ✓ |  |
| Peterson et al. (2020) | 15.4a | Painter et al. (2014)<br><u>Original data (113 stands)</u> | 2003-2012<br>2001-2016<sup>¶</sup> | ✓ | ✓ | ✓ | ✓ |

\* Authors sampled the five tallest young aspen within a stand, except for Halofsky et al. (2008) who sampled the three tallest young aspen.
<sup>†</sup>Included data from a random sample that provided no information on annual trends in height, and little or no information on annual trends in herbivory; the latter was limited to changes in browsing during 1997-1998 and 2011-2012 reported in Painter et al. (2014, 2015).
<sup>‡</sup>Did not include data for 2000 and 2003.
<sup>§</sup>Did not specify the data source for 2002.
<sup>¶</sup> Did not include data for 2015.

Although the first study that sampled only the tallest young aspen acknowledged that such “data are only representative of the first recovering aspen…and not an estimate of aspen population response across Yellowstone’s northern range” (Ripple & Beschta 2007), subsequent studies emphasized that the tallest young aspen represent a “leading edge” indicator of the future condition of the aspen population (e.g., Ripple & Beschta 2012; Painter *et al*. 2014; Beschta *et al*. 2018). Peterson et al. (2020) explained that sampling the tallest young aspen “allows investigators to document the occurrence of any young aspen exceeding the upper browse level of elk (i.e., ∼200 cm) in a given stand years before the average stem height attains this metric”.However, the fate of the tallest young aspen may not represent that of the average young aspen if the former are exposed to more favorable growing conditions than the latter. For example, if the tallest young aspen exceed the preferred browsing height, and therefore experience progressively less browsing and more height growth as they grow taller, they may realize a faster growth rate that leads to a level of overstory recruitment that is unattainable for the average young aspen. Thus, sampling the tallest young aspen may misrepresent the current and future aspen population response to wolf reintroduction.

We assessed the scope of this bias with data from E. Larsen’s long-term study of young aspen in northern YNP. These data reveal that the customary practice of sampling the tallest young aspen overestimated the potential for the typical young aspen to escape herbivory and reach tree height following wolf reintroduction, and thus exaggerated a popular example of a trophic cascade featured in many textbooks (e.g., Chapin *et al*. 2011; Boyce *et al*. 2012) and media (e.g., Morell 2007; Monbiot 2014). Our results highlight the importance of principled sampling for understanding trophic cascades following the loss and recovery of large predators.

## MATERIALS AND METHODS

### Study area

We measured young aspen in the portion of the northern Yellowstone elk winter range that lies within Yellowstone National Park (Fig. 1). This 995-km^2^ area is defined by low-elevation (2000-2600 m) grasslands and shrub steppes that fan out from the Yellowstone River and its tributaries near the Park’s northern border. A variety of ungulates spend winter in the area including elk, *Bison bison* (bison), *Odocoileus hemionus* (mule deer), and *Alces alces* (moose). Elk and bison were the most abundant ungulate in the area during winter (1,200-6,000 elk; 1,400-3,200 bison), and bison numbers exceeded those of elk beginning winter 2011-2012 (Tallian *et al*. 2017). Wolves were reintroduced to YNP in 1995-1997 (Bangs & Fritts 1996) and their annual distribution was concentrated in the study area (Cassidy *et al*. 2020) where they hunted mainly elk (Metz *et al*. 2020). Other elk predators included *Puma concolor* (cougar), *Ursus arctos* (brown bear), and *Ursus americanus* (black bear) (Barber-Meyer *et al*. 2008; Ruth *et al*. 2019). Elk that moved beyond the study area into adjacent areas of Montana were also hunted by humans (Peterson *et al*. 2020).

**Figure 1.**
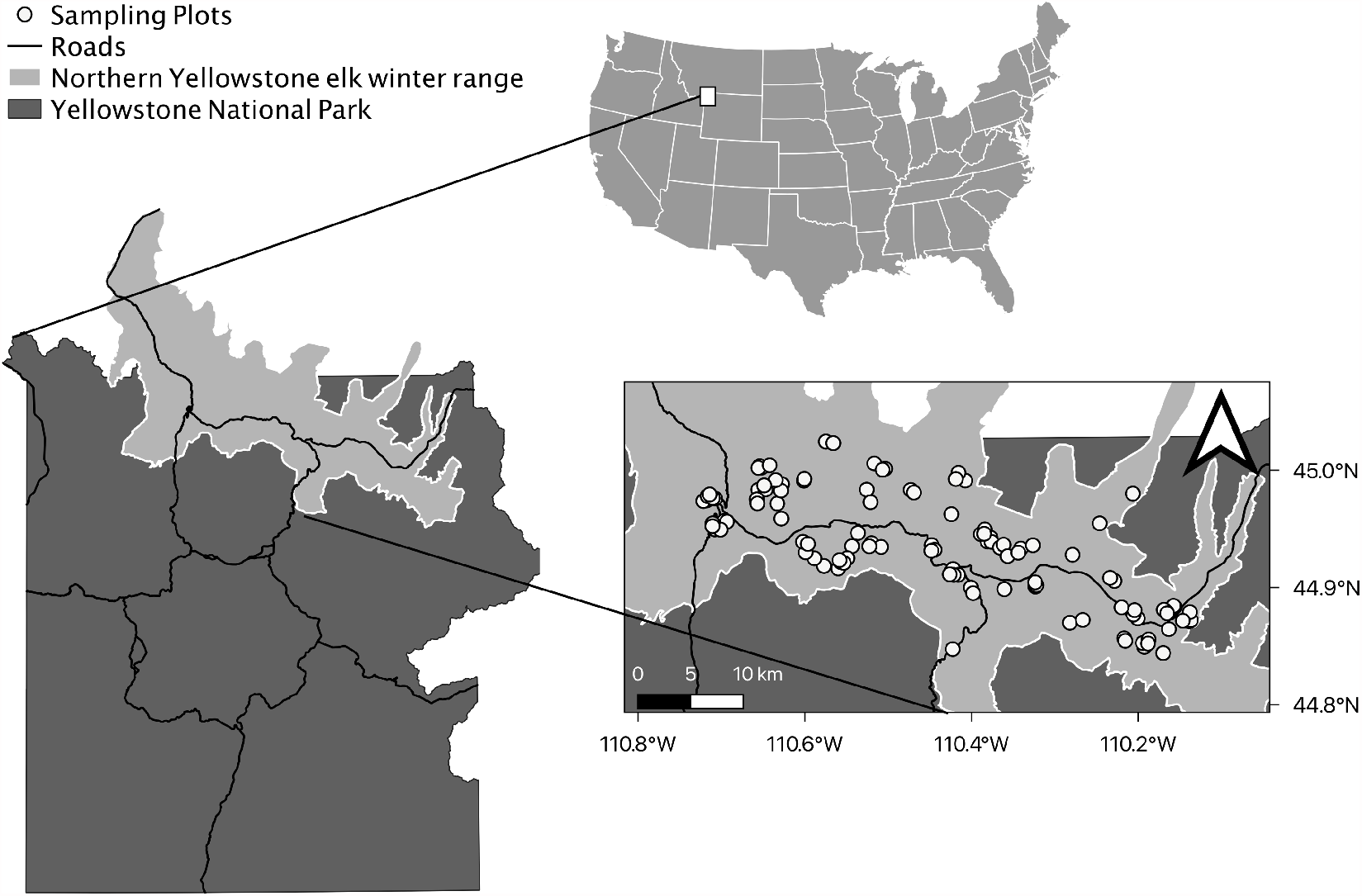
Locations of randomly sampled aspen stands in northern Yellowstone National Park. The northern Yellowstone elk winter range is the maximum distribution of the northern Yellowstone elk population during winter when elk often browse young aspen.

### Study population

Aspen is one of the few upland deciduous tree species in YNP and it is scattered across the study area in discrete stands on relatively moist mid-elevation benches, near streams, and along conifer forest/shrub steppe ecotones (Houston 1982). Aspen is a clonal species that mainly regenerates by root sprouting with occasional seedling establishment after disturbance. Root sprouting produces genetically identical trees from a common root system that may be substantially older than the age of the oldest tree, which rarely exceeds 150 years. Aspen requires moist soils and occurs mostly in areas with ≥ 38 cm of annual precipitation (Jones & DeByle 1985); the study area is near this lower limit (Larsen & Ripple 2003). Although aspen is a minor cover type in the arid portions of its range, it is a major source of biological diversity, providing habitat for numerous plants and animals (DeByle 1985; Mueggler 1985). Various ungulates eat the leaders and twigs of young stems, especially during winter. Persistent browsing of a stem’s leader prevents its growth to tree height, a process that has contributed to loss of overstory aspen in portions of western North America, including the study area (National Research Council 2002; Seager et al. 2013).

Historic landscape photographs suggest that overstory aspen covered ∼4-6% of the study area during 1880-1900 (Houston 1982; Meagher & Houston 1999), and aerial photographs indicate that aspen coverage decreased to ∼1% by the 1990s (Larsen & Ripple 2005). Pre-wolf reintroduction planning studies did not predict any cascading effects of wolves on aspen (reviewed in Yellowstone National Park 1997). Subsequent studies proposed that regeneration of overstory aspen depended on wolves reducing elk browsing (White *et al*. 1998; Ripple & Larsen 2000). The first evidence of substantial numbers of unprotected aspen reaching tree height since the early- to mid-20^th^ century (Larsen & Ripple 2003; Kauffman *et al*. 2010) occurred about a decade after wolf reintroduction (Ripple & Beschta 2007). See Appendix S1 in Supporting Information for additional context.

### Data collection

We measured browsing and height of young aspen in 113 plots distributed randomly across the study area (Fig. 1). Each plot was a 1 m × 20 m belt transect located randomly within an aspen stand that was itself randomly selected from an inventory of stands. The inventory was a list of 992 grid cells (240 m × 360 m) across the study area that contained at least one stand (Appendix S1). A “stand” was a group of tree-size aspen in which each tree was ≤ 30 m from every other tree. One hundred and thirteen grid cells were randomly selected from the inventory (∼11% of 992 cells), one stand was randomly selected from each cell, and one plot was established in each stand, with the start point and transect direction randomly determined (Ripple *et al*. 2001). We verified that each plot likely represented a genetically unique stand (Appendix S1).

We measured aspen at the end of the growing season (late July to September), focusing on plants ≤ 600 cm tall, which we termed “young aspen.” For each stand, we measured every young aspen rooted within a plot (‘random stems’), as well as each of the five tallest young aspen rooted anywhere within the stand (‘5T stems’). For all young aspen, we measured browsing (browsed or unbrowsed) and height of the leader stem (the tallest stem). A leader was ‘browsed’ if its growth from the previous growing season had been eaten. Most plots were measured nearly every year since 1999 (Ripple *et al*. 2001) and our analysis focused on data from 10 years (2007-2014, 2016-2017) in which sampled stands included measurements of random and 5T stems. Camera trap data indicate that elk were likely the primary ungulate species browsing young aspen in our plots during the study (Fig. S1).

### Data analysis

First, we built empirical distributions of browsing levels and stem heights to visualize how these characteristics differed between 5T and random stems within and across years (Appendix S1). Second, we modeled annual changes in browsing and stem height, and tested how these changes differed between 5T and random stems. Third, we modeled the influence of stem height on browsing to identify the preferred browsing height (PBH) and browse escape height (BEH). We used these results to assess the prevalence of non-preferred stems in samples of 5T and random stems, and to test how estimates of overstory recruitment differed between random and 5T stems.

#### Annual changes in browsing, stem height, and overstory recruitment

We combined measurements of 5T and random stems into one dataset of all stems (N = 18,623) across all years (N = 10 years) and used generalized linear mixed models (GLMMs) to test how the effect of year on browsing, height, and recruitment of stems differed by sampling method.

We treated the stem as the unit of analysis and used GLMMs with a Bernoulli distribution and a logit link to separately analyze the probability a stem was browsed (1 = browsed; 0 = not browsed) and recruited (1 = recruited; 0 = not recruited), and GLMMs with a gamma distribution and a log link to analyze stem height (cm), which took only non-negative values that were strongly right-skewed. A stem ‘recruited’ if it exceeded a presumed BEH of 200-cm or 300-cm. Year was an integer that ranged from 1 (2007) to 11 (2017) and sampling method was a dummy variable (1 = 5T sampling; 0 = random sampling). If 5T sampling estimates a faster annual decrease in browsing and faster annual increases in height and recruitment compared to random sampling, we expected (*i*) models with a year × method interaction fit the data better than models with only main effects for these variables, and (*ii*) the sign of this interaction was negative in the browsing model and positive in the height and recruitment models. We used likelihood ratio tests to compare models.

All GLMMs included a random intercept for stand identity, and GLMMs of browsing and stem height also included a random slope for year. We observed too few stems >200 cm or >300 cm to estimate a GLMM of recruitment with a random slope for year. The random intercept controlled for correlation among measurements of the same stand in multiple years and unmeasured stand-related effects including soil, water, and light conditions; the random slope permitted stand-specific annual trends in herbivory and stem height. We estimated average marginal effects (AMEs) from GLMMs to quantify and compare annual changes of 5T and random stems. AMEs describe the average effect of changes in explanatory variables on the change in a response variable and are useful for interpreting generalized linear models (Leeper 2021). We used statistics to test how the AMEs of year on browsing, stem height, and recruitment differed between 5T and random stems.

We calculated population-averaged fitted values from best-fit GLMMs by deriving marginal expectations of the responses averaged over the random effects but conditional on the observed variables. We refit year as a categorical factor and plotted the associated fitted values to illustrate the distribution of the underlying data after controlling for stand-level heterogeneity, and to assess the negative correlation between mean annual browsing and mean annual stem height.

Following previous studies (Painter *et al*. 2014, 2015, 2018; Beschta *et al*. 2016), we also estimated recruitment at the stand level as the percentage of sampled stands with stems taller than the presumed reach of elk. We calculated this separately for 5T and random stems as the annual percentage of sampled stands in which the median stem height exceeded 200 cm or 300 cm. Consistent with previous studies, recruitment estimates from 5T stems excluded stands that produced no young aspen.

#### Preferred browsing height and browse escape height

We modeled the effect of stem height on browsing to estimate PBH and BEH. We estimated separate GLMMs for 5T stems (N = 4,265) and random stems (N = 14,358), and included crossed random intercepts for stand identity and year to account for (*i*) correlation between measurements taken on the same stand in multiple years and on multiple stands in the same year, and (*ii*) unmeasured stand- and year-related effects.

We used piecewise linear splines to identify the PBH, which we defined as the height threshold beyond which browsing probability decreased with further height increase. We compared models with a single height threshold placed from 10-200 cm (first at 10 cm and then 1 cm intervals), a model with no height threshold, and an intercept-only model. We selected thresholds *a priori* based on the traditional assumption that stems taller than 200 cm escape browsing (Kay 1990). We constructed variables containing a linear spline for stem height so that the estimated coefficients measure the slopes of the segments before and after the threshold. We evaluated competing models using marginal likelihoods and information-theoretic statistics (AIC_c_; Burnham & Anderson 2002), and used AMEs of the best models to estimate how browsing probability changed with increasing stem height. We also used the best models to identify the BHE, which was the stem height at which browse probability was near zero.

## RESULTS

### Empirical distributions of browsing and stem height

Probability densities indicate that low browsing (≤ 20% stems browsed) and tall height (>100-cm) were more characteristic of 5T stems than of random stems, and that these differences increased from 2007 to 2017 (Fig. 2; Fig. S2). During this period, median height of 5T stems tracked increases in the 85-90^th^ percentile height of random stems, which increased 4-5 times faster than did the median height of random stems (Fig. S3).

**Figure 2.**
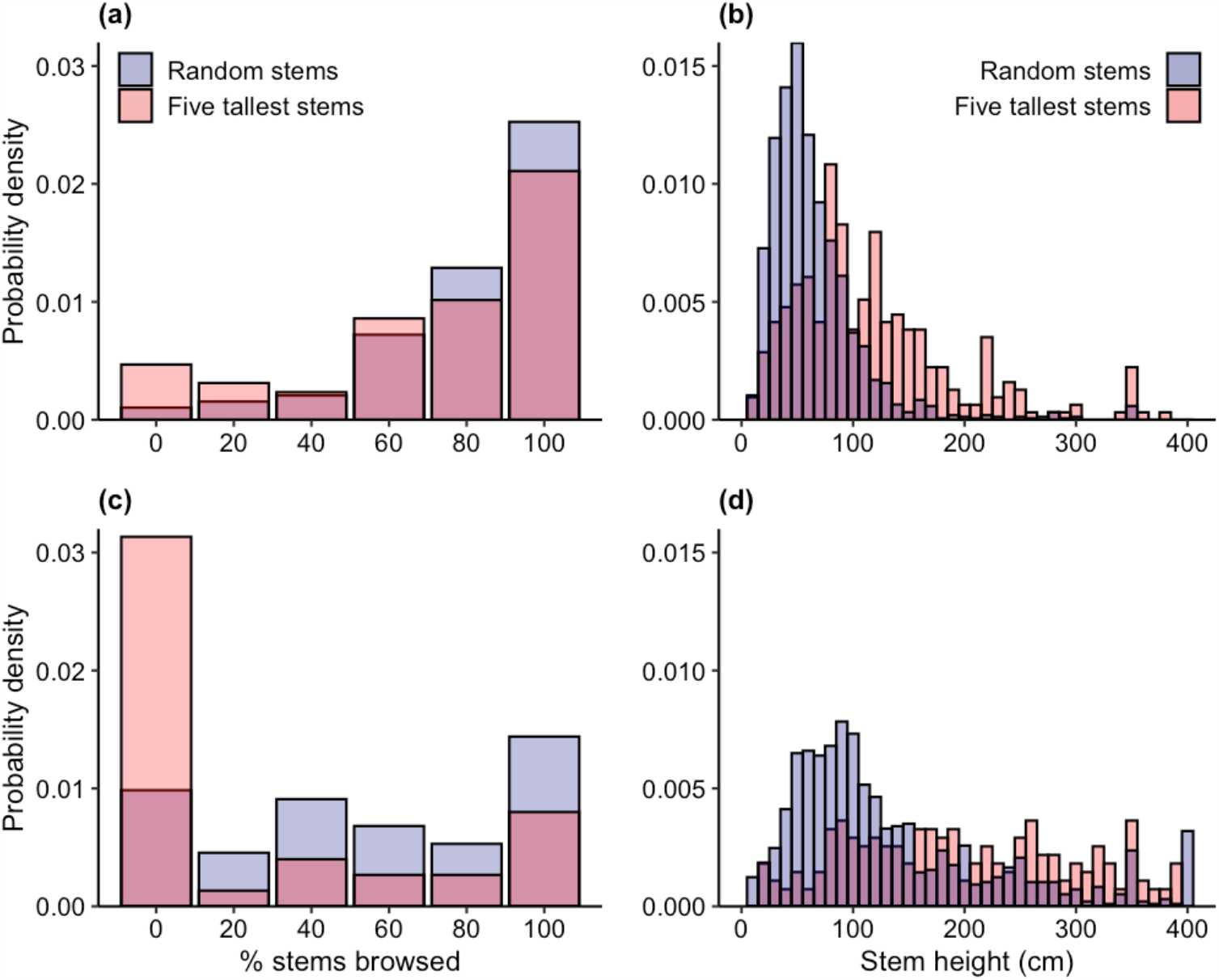
Probability densities of browsing (a, c) and height (b, d) of the five tallest young aspen and randomly sampled young aspen in northern Yellowstone National Park during the first and last years of the study (2007, 2017). Low browsing levels and tall heights were more characteristic of the five tallest young aspen throughout the study from 2007 (a, b) to 2017 (c, d). Probability densities for each year of the study are provided in Fig. S2.

### Annual changes in browsing and stem height

GLMMs of browsing and stem height with a year × method interaction fit the data better than did those with only main effects for these variables (browsing: 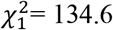, *P* < 0.001; height: 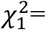47.2, *P* < 0.001). The interaction was negative in the browsing model (β = -0.18; 95% CI = -0.22, 221 -0.15; *P* < 0.001) and positive in the height model (β = 0.016; 95% CI = 0.012, 0.021; *P* < 0.001), which indicates that 5T sampling estimated a faster decrease in browsing and faster increase in height compared to random sampling (Fig. 3a, 3b). Specifically, browsing decreased 3.9 percentage points·year^-1^ (95% CI = 3.2, 4.6) for 5T stems versus 1.8 percentage points·year^-1^ (95% CI = 1.2, 2.5) for random stems (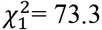, *P* < 0.001). Height increased 19.8 cm·year^-1^(95% CI = 15.8, 23.9) for 5T stems versus 7.7 cm·year^-1^ (95% CI = 6.0, 9.4) for random stems (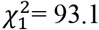, *P* < 0.001). The negative correlation between estimates of mean annual browsing (points in Fig. 3a) and mean annual height (points in Fig. 3b) was 47% stronger for 5T stems (Pearson’s correlation coefficient, *r* = -0.97; *P <* 0.001) compared to random stems (*r* = -0.60; *P* 230 = 0.07; Fig. 3c).

**Figure 3.**
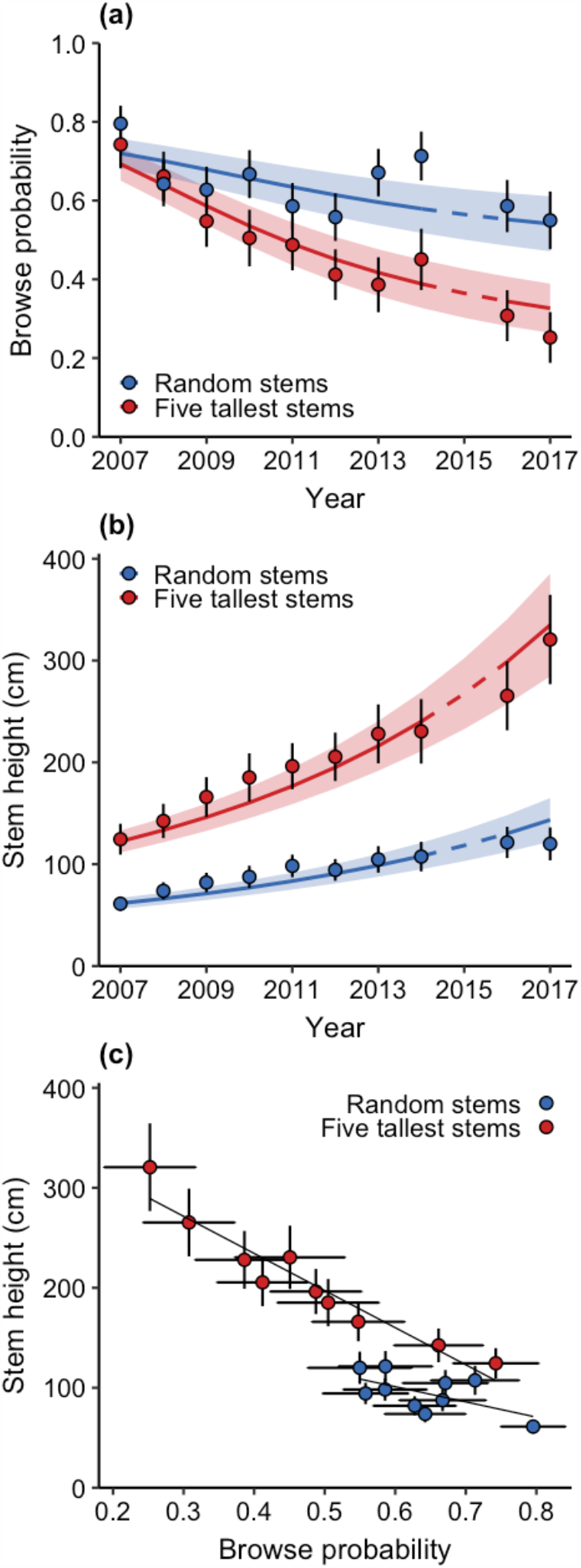
Effects of sampling method on estimated annual trends in browsing and height of young aspen in northern Yellowstone National Park, 2007-2017. Relative to a random sample, a nonrandom sample of the five tallest stems estimated a faster annual decrease in browsing (a), faster annual increase in stem height (b), and stronger negative correlation between browsing and stem height (c). Results in (a) and (b) are population-averaged fitted values and associated 95% confidence intervals from best-fit GLMMs of the interactive effect of year and sampling method on browse probability and stem height with year modeled as a continuous (lines) or categorical (points) effect. Results in (c) are the relationships between the categorical fitted values in (a) and (b), with lines estimated from simple linear regressions. No data were collected in 2015.

### Preferred browsing height and browse escape height

The best-fit GLMMs (ΔAIC_c_ = 0) indicate that the PBH was 132 cm for 5T stems and 122 cm for random stems (Fig. 4a; Tables S1, S2). Below these heights, each 10 cm increase in height increased browsing by 0.4 percentage points for 5T stems (95% CI = -0.3, 1.0; *P* = 0.26) and 0.5 percentage points for random stems (95% CI = 0.1, 0.8; *P* = 0.005). Above these heights, each 10 cm increase in height decreased browsing by 3.1 percentage points for 5T stems (95% CI = 2.9, 3.2; *P* < 0.001) and 3.5 percentage points for random stems (95% CI = 3.2, 3.7; *P* < 0.001). Stems exceeding the PBH were 1.6-5.1 times (mean ± SE = 2.8 ± 0.40) more prevalent in the sample of 5T stems than in the sample of random stems (Fig. 4b). The best-fit GLMMs indicate that browsing of stems >200 cm was as high as 0.45 (95% CI = 0.40, 0.51) for 5T stems and 0.35 (95% CI = 0.28, 0.42) for random stems. Browsing was negligible (< 0.07) only after stems exceeded ∼300 cm (Fig. 4a).

**Figure 4.**
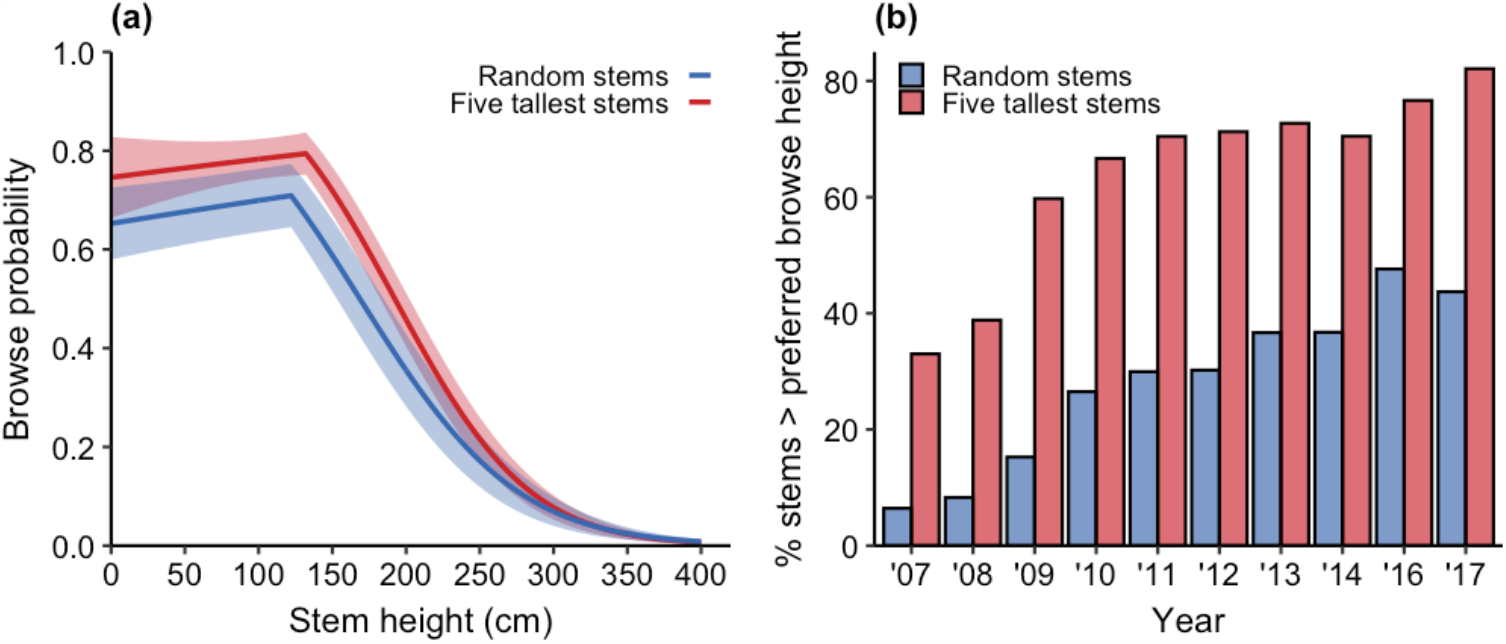
Effects of stem height on the probability a young aspen stem was browsed (a), and the annual percentage of stems in the sample of young aspen that exceeded the preferred browsing height of 132 cm (five tallest stems) or 122 cm (random stems) (b) in northern Yellowstone National Park, 2007-2017. Lines in (a) are population-averaged fitted values and associated 95% confidence intervals from best-fit GLMMs estimated separately for five tallest stems (Table S1) and random stems (Table S2). Bars in (b) are percentages of the total annual sample size – pooled across plots – of five tallest stems (N=317-518 stems·year^-1^) and random stems (N=1027-1748 stems·year^-1^). No data were collected in 2015.

### Overstory recruitment

GLMMs of the probability that a stem exceeded 200 cm or 300 cm with a year × method interaction fit the data better than did those with only main effects for these variables (200 cm: 248 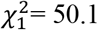, *P* < 0.001; 300 cm: 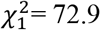, *P* < 0.001). The positive sign of the interaction (200 cm: β = 0.23; 95% CI = 0.17, 0.30; *P* < 0.001; 300 cm: β = 0.40; 95% CI = 0.30, 0.49; *P* < 0.001) indicates that 5T sampling estimated a faster increase in recruitment compared to random sampling (Fig. 5a, 5b). Recruitment of stems >200 cm increased 4.6 percentage points·year^-1^(95% CI = 3.9, 4.2) for 5T stems versus 1.5 percentage points·year^-1^ (95% CI = 1.1, 1.9) for random stems (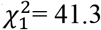, *P*<0.001), and recruitment of stems >300 cm increased 3.1 percentage points·year^-1^ (95% CI = 2.5, 3.6) for 5T stems versus 0.49 percentage points·year^-1^ (95% CI = 0.32, 0.67) for random stems (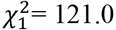, *P*<0.001).

**Figure 5.**
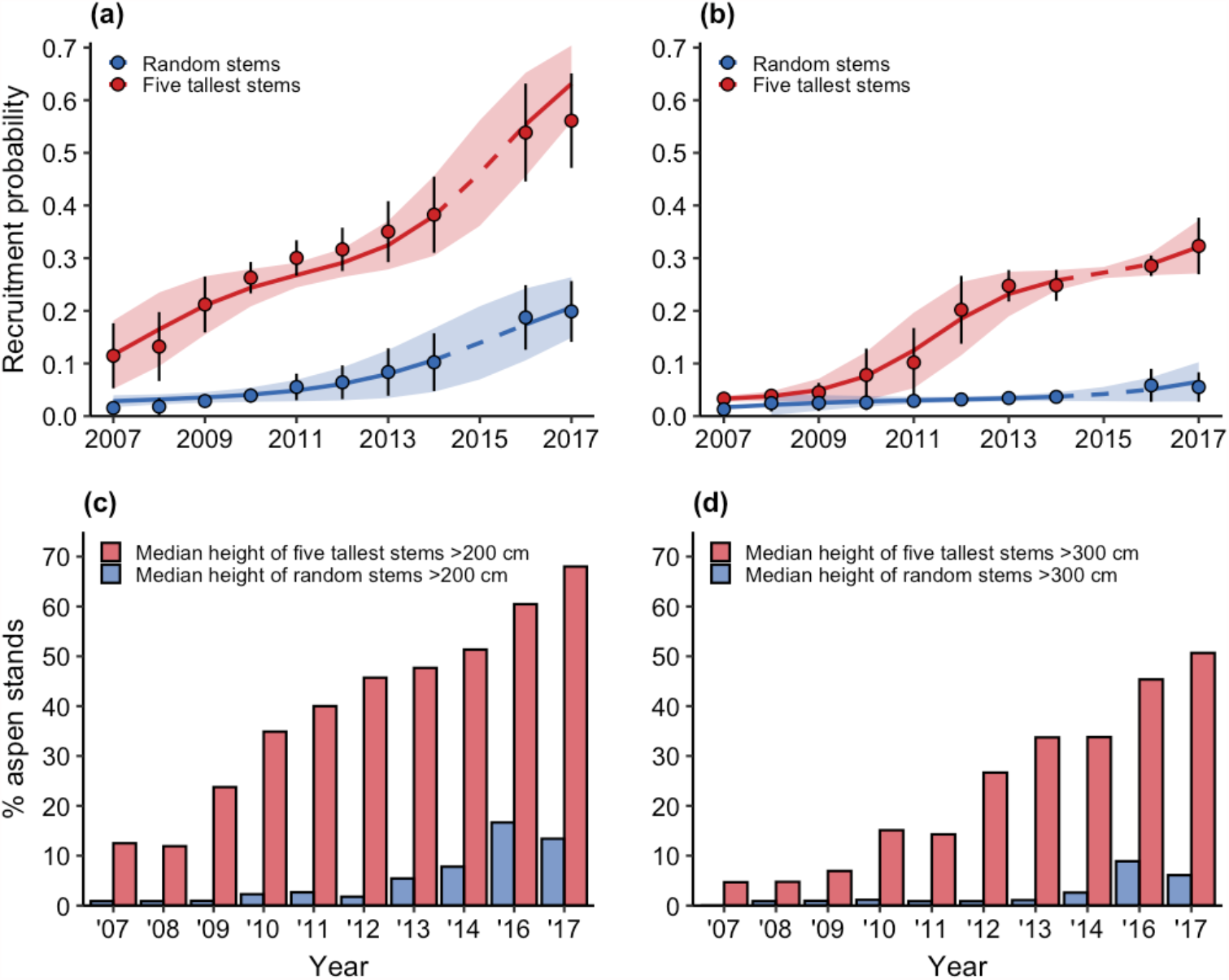
Effects of sampling method and browse-escape height assumption on stem-level (a, b) and stand-level (c, d) estimates of annual trends in overstory aspen recruitment in northern Yellowstone National Park, 2007-2017. Sampling the five tallest aspen under the assumption that stems taller than 200 cm escaped browsing and joined the overstory estimated rapid annual increases in overstory recruitment (a, c), whereas random sampling all young aspen under the assumption that stems taller than 300 cm escaped browsing and joined the overstory estimated relatively slow annual increases in overstory recruitment (b, d). Results in (a) and (b) are population-averaged fitted values and associated 95% confidence intervals from best-fit GLMMs of the interactive effect of year and sampling method on the probability that a stem exceeded 200 cm (a) or 300 cm (b) with year modeled as a continuous (lines) or categorical (points) effect. Bars in (c) and (d) are percentages of the total annual sample of aspen stands in which the median height of the five tallest stems and randomly sampled stems exceeded a presumed browse-escape height of 200 cm or 300 cm. No stands had a median height of random stems >300 cm in 2007 and no data were collected in 2015.

Stand-level assessments indicate that 5T sampling estimated a total increase in overstory recruitment from 2007 to 2017 that was 4-8 times greater than that estimated by random sampling. The annual percentage of stands with a median stem height greater than 200 cm (300 cm) increased from 13-68% (5-51%) for 5T sampling versus 1-13% (0-6%) for random sampling, and in any given year, the percentage estimated from 5T sampling was 4-26 times (5-31 times) greater than that estimated by random sampling (Fig. 5c, 5d). Random sampling revealed that young aspen were annually absent in 2-9% of stands and 11-19% of plots within stands (Fig. S4, S5). Every stand produced young aspen during at least one year of the study, whereas 7% of plots consistently produced no young aspen.

## DISCUSSION

To the extent that annual decreases in browsing and annual increases in height of woody deciduous plants in northern Yellowstone National Park reflect the cascading effects of reintroduced wolves (Beyer *et al*. 2007; Beschta & Ripple 2016; Beschta *et al*. 2018), our results show that an accepted practice of nonrandom sampling – one featured in a dozen peer-reviewed publications over more than a decade (Table 1) – exaggerated the aspen population response to wolf reintroduction. Sampling only the five tallest young aspen within a stand estimated annual changes in browsing, stem height, and overstory recruitment that were significantly faster than those estimated by random sampling of all young aspen within a stand (Fig. 3a-b, Fig. 5). We suggest that 5T sampling exaggerated the aspen response for the following three reasons.

First, 5T sampling favored stems that were taller than the preferred browsing height (PBH) of elk (Fig. 4). Stems taller than the PBH likely grew faster than stems shorter than the PBH because the former were browsed at a decreasingly low rate as they grew taller, whereas the latter were browsed at an increasingly high rate as they grew taller (Fig. 4a). Stems taller than the PBH may have also allocated relatively fewer resources to defense chemistry expression as browsing decreased (Rhodes *et al*. 2017), which would have further accelerated their height growth (Lindroth & St. Clair 2013).

The PBH that we estimated (122-132 cm) is similar to the PBH of elk and comparable cervids (red deer, *Cervus elaphus*) in other systems (Rounds 1979; Motta 2003; Renaud *et al*. 2003; Konôpka *et al*. 2018). Experiments indicate that the PBH equals ungulate shoulder height, and that ungulates prefer to browse at this height because it minimizes neck angle which in turn maximizes foraging efficiency (Renaud *et al*. 2003). Although published shoulder height estimates for Yellowstone elk are lacking, the literature suggests a range of 119-168 cm (Murie 1951; Blood & Lovaas 1966; Blood & Smith 1984; Hudson & Haigh 2002). Camera trap photos of elk browsing aspen in our plots is consistent with the hypothesis that elk prefer to browse at the level of their shoulder height (Fig. S6).

Our PBH estimate is also similar to (or less than) previous estimates of mean height of the tallest young aspen (Table 1), which implies that these studies also measured a large number of stems taller than the PBH. These estimates refer to the 1-2 years in which field sampling occurred as well as to preceding years that lacked field sampling. Estimates for the latter were retrospectively inferred using plant architecture methods. Although such methods might not be useful when herbivory is intense (Ripple & Beschta 2007), the prevailing assumption is that such estimates are accurate indicators of browsing and stem height across decadal time scales. To the extent that this assumption is valid, our results suggest that reported trends in browsing and stem height based on the tallest young aspen (Table 1) do not represent trends in the aspen population at large because they too were biased toward stems that were taller than the PBH.

A challenge highlighted by the PBH is that stem height is both a cause and an effect of reduced browsing. Thus, the negative correlation between browsing and stem height (e.g., Fig. 3c) that some consider critical evidence of a trophic cascade (Beyer *et al*. 2007; Beschta & Ripple 2016; Beschta *et al*. 2018) is not an exclusive indicator of browsing suppressing stem height. Rather, the negative correlation also indicates that stem height suppresses browsing, which reflects previous findings that factors besides browsing control stem height (Romme *et al*. 1995). This relationship helps explain why the negative correlation between browsing and stem height was stronger for 5T stems compared to random stems (Fig. 3c): the stronger negative correlation was consistent with the strong negative effect of stem height on browsing that most 5T stems experienced given most were taller than the PBH. Similarly, the weak negative correlation for random stems was at least partially due the countervailing positive effect of stem height on browsing that most of these stems experienced since most were shorter than the PBH (Fig. 4). These results suggest that a negative correlation between browsing and stem height is not reliable evidence of a trophic cascade because it does not represent an unambiguous causal link between reduced browsing and increased stem height.

Second, the tallest young aspen were most likely exposed to the best growing conditions. Tall stature is itself an indicator of a locally productive resource environment because only stems exposed to sufficient sunlight, moisture, and soil quality have the capacity to grow tall (Brown *et al*. 2006; Hansen *et al*. 2016). Thus, more productive resource conditions may have contributed to the faster annual height growth and overstory recruitment of 5T stems (Fig. 3b, Fig. 5). Faster height growth could have in turn contributed to the faster decrease in herbivory (Fig. 3a). It is also possible that better resource conditions permitted 5T stems to respond more rapidly to a reduction in herbivory (Fig. 3c).

Third, 5T sampling did not detect the absence of aspen regeneration. By definition, 5T sampling measures only stands and locations within stands that produce young aspen. Stands and locations without young aspen are not sampled and therefore excluded from stand-level estimates of overstory recruitment calculated as the percentage of sampled stands with stems taller than the presumed reach of elk (e.g., Fig. 7b in Painter *et al*. 2014, Fig. 2b in 2015; Fig. 5d in Beschta *et al*. 2016). This omission likely inflates stand-level estimates of overstory recruitment given our finding that young aspen were annually absent in 2-9% of stands and 11-19% of plots within stands (Fig. S4). Overestimating recruitment because the absence of regeneration is undocumented in poorly recruiting stands represents a form of visibility bias that has received little attention in discussions about the cascading effects of wolves on aspen.

A separate source of bias that further inflates stand- and stem-level estimates of overstory recruitment is the traditional assumption that stems taller than 200 cm escape herbivory. Our analysis indicates that escape from herbivory is not certain until stems exceed approximately 400 cm (Fig. 4a). This suggests that previous estimates of overstory recruitment that assume a browse escape height of 200 cm (Painter *et al*. 2014, 2015, 2018; Beschta *et al*. 2016) are best interpreted as maximum estimates.

Our results do not support the hypothesis that the tallest young aspen represent a “leading edge” indicator of a “broader shift in plant community dynamics for northern [YNP] aspen stands” (Beschta *et al*. 2018). Contrary to the expectation that characteristics of the average young aspen should resemble those of the tallest young aspen over time (Peterson *et al*. 2020), we found that mean levels of herbivory, height, and recruitment of random stems increasingly differed from those of 5T stems (Fig. 3a-b; Fig. 5). Trends in the height distributions of 5T stems relative to those of random stems indicate that this widening gap was because 5T stems represented a minority of young aspen that substantially outperformed the majority of young aspen. Specifically, height increase of 5T stems tracked the height increase of the 85-90^th^percentile of random stems, which was substantially faster than the height increase of more than half of random stems (Fig. S3). Assuming our sample of random stems was representative of the population at large, our results suggest that the dynamics of 5T stems represent those of the top-performing ∼10-15% of young aspen.

The general lesson from our study is that apparently minor deviations from principled sampling can lead to major misunderstandings about the strength of trophic cascades in response to large predator recovery. Had we sampled only the tallest young aspen, we would have concluded, incorrectly, that wolf reintroduction was associated with a strong regenerative response of aspen. Instead, our random sampling design, which included a more representative sample of aspen stands and stems, indicated that wolf reintroduction was associated with a more limited regenerative response that did not reverse the deterioration of all aspen stands (e.g., Fig. S5). A limited aspen response is consistent with (*i*) documented losses of other aspen stands in the study area since wolf reintroduction (Fig. S7; Beschta *et al*. 2020), (*ii*) evidence that wolves had weak and/or inconsistent effects on elk foraging behavior (Laundré *et al*. 2001; Childress & Lung 2003; Wolff & Van Horn 2003) and habitat selection (Kohl *et al*. 2018, 2019; Cusack *et al*. 2019), and weak to moderate effects on elk population density (Vucetich *et al*. 2005; Peterson *et al*. 2014; Macnulty *et al*. 2020), and (*iii*) theory that predicts weak cascading effects in food webs like Yellowstone that are resource-limited, spatially heterogeneous, and reticulated (Ford & Goheen 2015). We also note that climate change may have contributed to the limited aspen response that we documented given that many of our plots occurred in or adjacent to areas projected to become unsuitable for aspen due to anthropogenic climate forcing (Piekielek *et al*. 371 2015; Fig. S8).

Understanding how ecosystems respond to the loss and/or addition of large predators is vital to resolving broader debates about the forces that structure food webs, determine species abundance, and deliver ecosystem services (Estes *et al*. 2011; Dobson 2014; Ripple *et al*. 2014; Atkins *et al*. 2019). Our study, which focused on a textbook example of large predator extirpation and reintroduction, demonstrates how apparently harmless deviations from principled sampling can distort this understanding. Nonrandom sampling overestimated the strength of a hypothesized trophic cascade in the system we studied, but it may underestimate cascading effects in other systems. In observational studies that lack control, randomization is one of the few available protections against unreliable inferences and the misguided policy and management decisions they may inspire. Growing concerns about the reliability of research findings in ecology (e.g., Fraser *et al*. 2002; Fidler *et al*. 2017; O’Grady 2020) emphasize the importance of principled sampling in studies of trophic cascades and other ecological phenomena.

## ACKNOWLEDGEMENTS

We thank R. Renkin and J. Klaptosky for administrative and field assistance. We also thank J. Brodie and N. Piekielek for sharing important supplemental data. We acknowledge support from the U.S. National Science Foundation (DGE-1633756), University of Wyoming-National Park Service Small Grant Program (1003867-USU), Yellowstone National Park, Utah State University Ecology Center, and the S.J. and Jessie E. Quinney Doctoral Fellowship. We thank Dr. William Ripple for serving as principal investigator on the scientific research and collecting permits issued by Yellowstone National Park that authorized aspen data collection during 1999-

## Authorship

E.M.B., E.J.L., D.R.M. conceived and designed the study; E.J.L. collected most of the data; E.M.B. and D.R.M. analyzed the data and wrote the manuscript with input from E.J.L.

## Data availability

Data are archived in Dryad (##).

