## Appendix S1 for "Sampling bias exaggerates a textbook example of a trophic cascade"

<u>Contents</u>

Appendix S1 Supplementary Materials, Methods, and Results

- Study population
- Aspen stand inventory
- Inter-plot genetic relatedness
- Species identity of ungulate browsers
- Empirical distributions of browsing and stem height
- Projections of aspen habitat suitability

References

Figure S1-S8

Table S1-S2

### Appendix S1 Supplementary Materials, Methods and Results

*Study population*

Poor regeneration of overstory aspen was noted in the study area (northern Yellowstone National Park) as early as the 1920s and attributed to herbivory from *Castor canadensis* (beaver; Warren 1926) and *Cervus canadensis* (elk; Rush 1932; Grimm 1939). As beaver abundance declined (Smith & Tyers 2012) and elk abundance remained high (Houston 1982), subsequent observers emphasized the role of elk herbivory in preventing aspen regeneration, with some considering it a proximal factor, secondary to fire suppression and climate variation (Houston 1982; Yellowstone National Park 1997; Singer *et al.* 1998), and others judging it an ultimate factor (Kay 1990; National Research Council 2002; Wagner 2006). Romme et al. (1995) concluded that the dearth of aspen regeneration was due to multiple factors including fire suppression, drying climate, and high levels of elk herbivory possibly linked to the extirpation of wolves.

*Aspen stand inventory*

Sampling plots were set randomly within an aspen stand that was itself randomly selected from an inventory of aspen stands. The inventory was a list of 992 landscape grid cells (240 m × 360 m) across the study area that contained at least one aspen stand. Aspen stands were identified from 83 color infrared aerial photographs (1:24,000) taken in October 1988. To identify stands, a 1.0 cm × 1.5 cm grid of 96 cells (240 m × 360 m ground dimensions) was overlain on each photograph, and a scanning stereoscope was used to identify all cells containing at least one aspen stand. In these aerial photographs, aspen were identifiable as white crowns, whereas conifers appeared as red crowns (Larsen & Ripple 2003).

*Inter-plot genetic relatedness*

We tested whether our plots represented genetically independent samples given that aspen clones can sprout genetically identical stems up to 40-m away (Rogers *et al.* 2020). To do so, we genotyped a sample of young aspen collected from a random subset of 59 plots (52% of total plots) at the end of the 2018 growing season.

We collected a single leaf from 1-4 young aspen in each plot (N = 122 young aspen). We sampled at least two young aspen in most plots (N = 54 plots), each rooted on opposite ends of the 20-m transect that defined each plot. Of these plots, seven included one additional young aspen sampled at the transect midpoint (10-m), and one included two additional young aspen sampled at 5-m and 15-m. In the five remaining plots, we sampled only a single young aspen, rooted at the transect start point, because young aspen were scarce in these plots. Each sampled leaf, which measured about 4.5-cm$^2$, was placed in a separate paper coin envelope and stored in an open container of silica gel to permit proper drying.

Genetic analyses were conducted at the Utah State University Molecular Ecology Laboratory. Each leaf sample was genotyped at 12 microsatellite loci, which provided sufficient resolution to determine genetic relatedness of plots. We calculated the Lynch-Ritland estimator of relatedness (Lynch & Ritland 1999) between all pairs of samples across plots, such that a relatedness coefficient of 1.0 indicated genetically identical young aspen. No pair met this criterion, consistent with our assumption that plots represented genetically unique aspen stands.

*Species identity of ungulate browsers*

We used camera trap photos collected in winters 2008-2009 (Brodie *et al.* 2012), 2018-2019, and 2019-2020 (this study) to assess the potential species identity of ungulates that browsed young aspen in our sample of stands (N = 113) in winter (01November-30April) during the focal years of our data analysis (2007-2014, 2016-2017). This assessment is approximate because cameras were either deployed in non-sampled stands during focal years (Brodie *et al.* 2012) or in sampled stands after focal years (this study). Camera data from Brodie et al. (2012) is applicable because the spatial distribution of their cameras approximated that of our sampled stands, with one camera overlapping (Plot 16, 44.959839° N, 110.692508° W) and all others within 20 m – 4 km of one of our stands (mean ± SE = 1.1 ± 0.25 km; Fig. S1a). Together, these data provide the best available information about the potential species identity of ungulates that browsed young aspen in our sampled stands during our period of interest.

In winter 2008-2009, Brodie et al. (2012) collected photos from 19 Reconyx RM45 cameras in each of 19 randomly selected aspen stands distributed across a snowpack gradient. Each camera was active 24 hours/day for 1-180 winter days (151.0 ± 10.8 days) (J. Brodie, personal comm.). We collected photos from a random subset of 20 stands in winter 2018-2019 using Bushnell Trail Cam HD and Moultrie cameras, and 13 stands in winter 2019-2020 using Bushnell Trail Cam HD cameras. Nine of the stands were monitored across both seasons and 15 were monitored for a single season, resulting in a total of 24 monitored stands. Cameras were deployed directly facing each plot, and were active 24 hours/day for 7-180 days (155.0 ± 9.54 days) in winter 2018-2019 and 172-180 days (179.0 ± 0.62 days) in winter 2019-2020.

To assess the identity of ungulates that browsed young aspen in our sampled stands across species and winters, we first calculated the relative abundance of each ungulate species sighted by each camera each winter. For both datasets (2008-2009 and 2018-2020), we defined a camera capture record as an independent sighting if it occurred at least 10 minutes after the previous record of an ungulate of the same species at the same camera (e.g., two photos of elk < 1 minute apart were assumed to be the same animal and classified as a single sighting). We processed photos manually or with the software Camelot (Hendry & Mann 2018), and for each independent sighting we determined species identity and number of individuals present. For each camera in each winter (2008-2009, 2018-2019, and 2019-2020), we calculated the relative daily abundance of each species as the total number of individuals sighted by the camera divided by the number of days the camera was active.

Second, we pooled sightings of ungulates engaged in the act of browsing and calculated the percentage of such "browse sightings" that featured each ungulate species. We defined "browsing" as an animal with its nose/mouth touching or reaching toward a young aspen. Data for this analysis were only available for winters 2018-2019 and 2019-2020.

Elk were the most numerous ungulate at monitored aspen stands near the start of the study in 2008-2009 (0.70 ± 0.23 elk/day, 0.19 ± 0.06 bison/day, and 0.04 ± 0.02 other ungulates/day; Fig. S1b) and were about as numerous as bison after the study in 2018-2020 (0.08-0.15 elk/day, 0.11-0.15 bison/day; Fig. S1b). Although the total number of bison wintering in the study area increased during our study (Tallian *et al.* 2017), the relative abundance of bison sighted by cameras at aspen stands was largely unchanged (Fig. S1b). The relative abundance of other ungulates (moose, mule deer, and pronghorn) was consistently low across the study period (Fig. S1b). Though these other ungulates are known to browse aspen (Stevens 1970; Singer & Norland 1994), elk were 19-30 times more abundant than these species after the study, which suggests that other ungulates contributed little to browsing. This conclusion is supported by photographic evidence: of the 29 browse sightings, 86% featured elk (N = 25), 10% featured moose (N = 3), and 4% featured mule deer (N = 1, Fig. S1c). There were no browse sightings of bison, which is consistent with earlier findings that aspen is a negligible portion of the diet of Yellowstone bison (Singer & Norland 1994). Thus, while relative abundance of elk and bison at our monitored stands was similar during winters 2018-2019 and 2019-2020, our browse sighting data indicate that elk were the dominant winter browser of aspen.

*Empirical distributions of browsing and stem height*

We calculated separate probability densities of browsing and height of random stems and 5T stems for each year of the study using kernel density estimation with a Gaussian distribution scaled to integrate to one (Fig S2). Individual stands and plots were the units of analysis in the probability densities of browsing, with stands and plots pertaining to 5T stems and random stems, respectively. Browsing of 5T stems equaled the percentage of the five tallest stems within a stand that were browsed, providing one of six possible values: 0%, 20%, 40%, 60%, 80%, and 100%. Browsing of random stems was the percentage of young stems within a plot that were browsed, and we rounded these values to the nearest of the above percentages to enable comparison with 5T stems. Individual stem was the unit of analysis in the probability densities of height, with individual heights pooled across stands and plots for each year.

*Projections of aspen habitat suitability*

Climate change has already reduced aspen occupancy across the western United States (Rehfeldt *et al.* 2009; Worrall *et al.* 2013), and this downward trend is expected to continue (Piekielek *et al.* 2015, 2016). Piekielek et al. (2015) assessed the fate of aspen in the Greater Yellowstone Ecosystem – including our study area – under representative concentration pathway (RCP) 8.5, which is consistent with increases in atmospheric greenhouse gases at present rates. Using aspen presence/absence data from the U.S. Forest Service Forest Inventory and Analysis (FIA) program, Piekielek et al. (2015) modeled current aspen distribution based on August water deficit, April snowpack, June soil moisture, rock volume, and percent sand in soil at a 30 arcsecond spatial resolution (~1 km). They then used 9 different global climate models under RCP 8.5 to project aspen habitat suitability in 2025, 2055, and 2085 (N. Piekielek, personal comm.). We plotted our sampling plots on these projections to assess the extent that sampled aspen stands were vulnerable to current and future climate change.

Projections were available for only those areas sampled by the FIA. Of our 113 sampling plots, 44 overlapped areas sampled by the FIA (Fig. S8). Of these 44 plots, all are in currently suitable aspen habitat that is projected to become unsuitable within the next 4-64 years. Eighteen plots are projected to be unsuitable by 2025, 25 plots are projected to be unsuitable by 2055, and one is projected to be unsuitable by 2085. No plots are in habitat that is projected to become suitable (Fig. S8). Based on these projections, we expect future reductions in aspen overstory recruitment in our sampling plots despite the influence of wolves and other predators on elk population dynamics.

**Appendix S1**: Supplementary materials for "Sampling bias exaggerates a textbook example of a trophic cascade"

**Figure S1.** Ungulate winter use of aspen stands in northern Yellowstone National Park, 2008-2009 and 2018-2020. (a) Map of camera locations from Brodie et al.'s (2012) study (green: winter 2008-2009), and this study (blue: winters 2018-2019 and 2019-2020). Open circles are the 113 sampling plots from this study. (b) Mean number (± SE) of individuals sighted per day across cameras each winter. Other ungulates = moose, mule deer, whitetail deer, and pronghorn. (c) Percentage of sightings of ungulates browsing young aspen by species. There were no photos of bison browsing young aspen. Data in (c) are from winters 2018-2019 and 2019-2020.

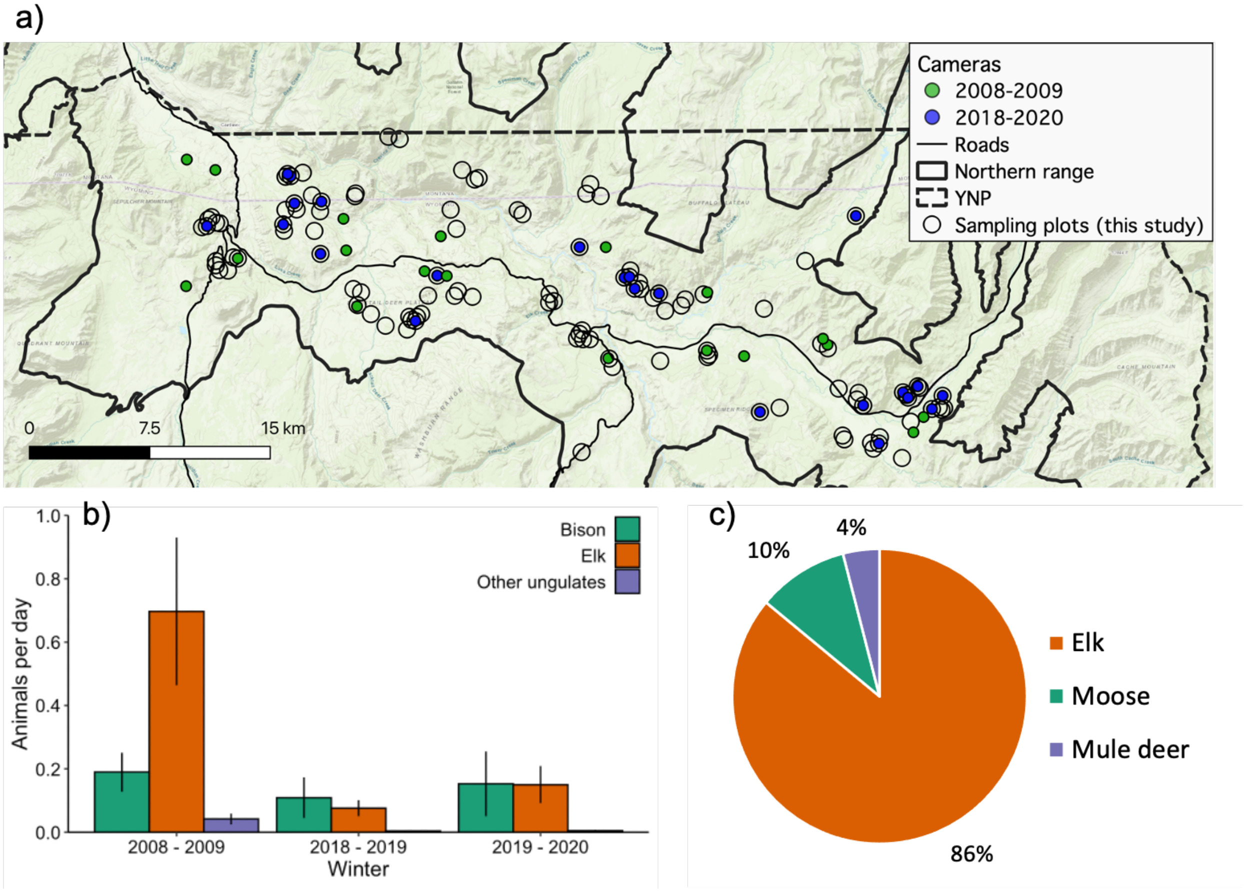

**Figure S2**. Annual probability densities of browsing (left panels) and height (right panels) of the five tallest young aspen and randomly sampled young aspen in northern Yellowstone National Park, 2007-2017. Data were not collected in 2015.

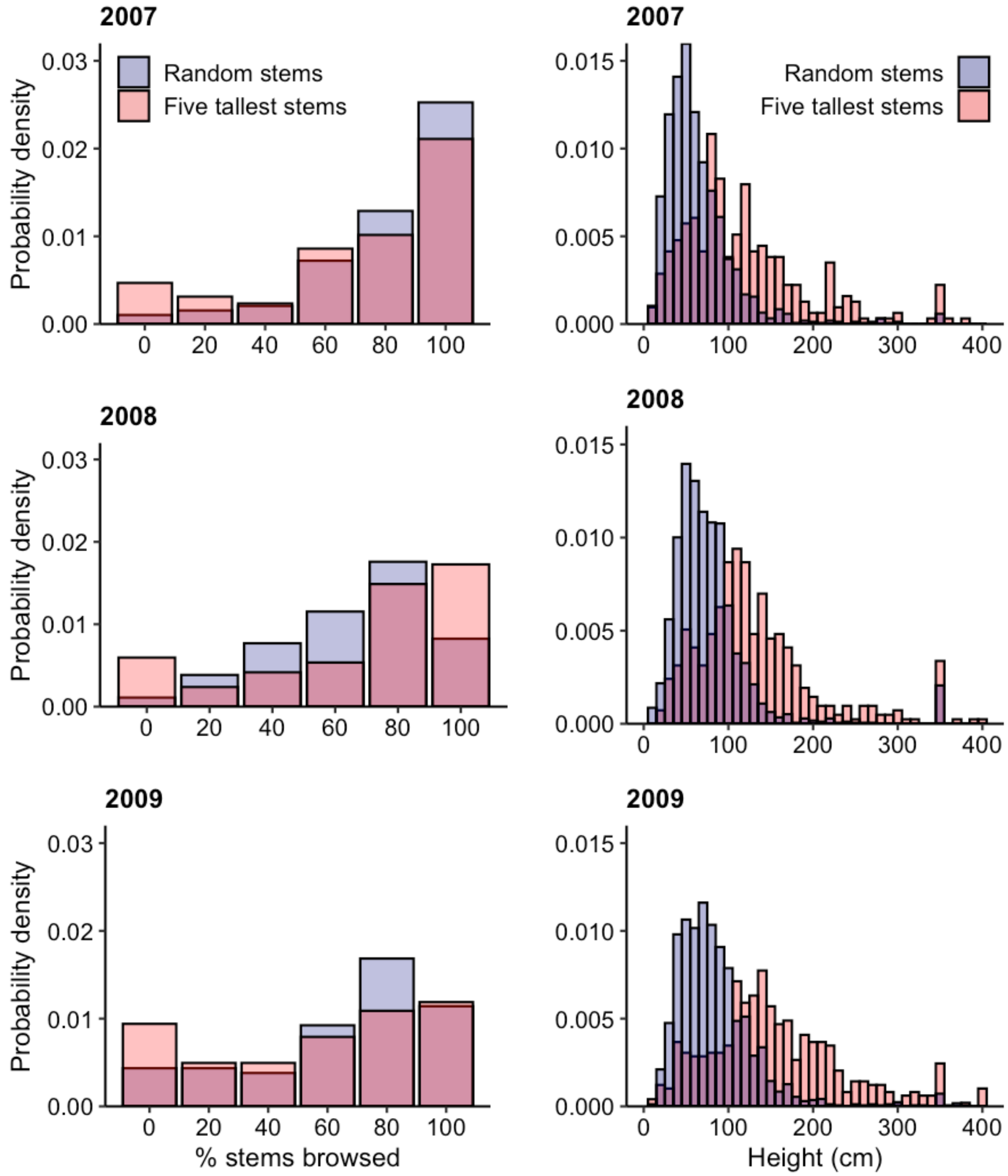

### Figure S2 continued

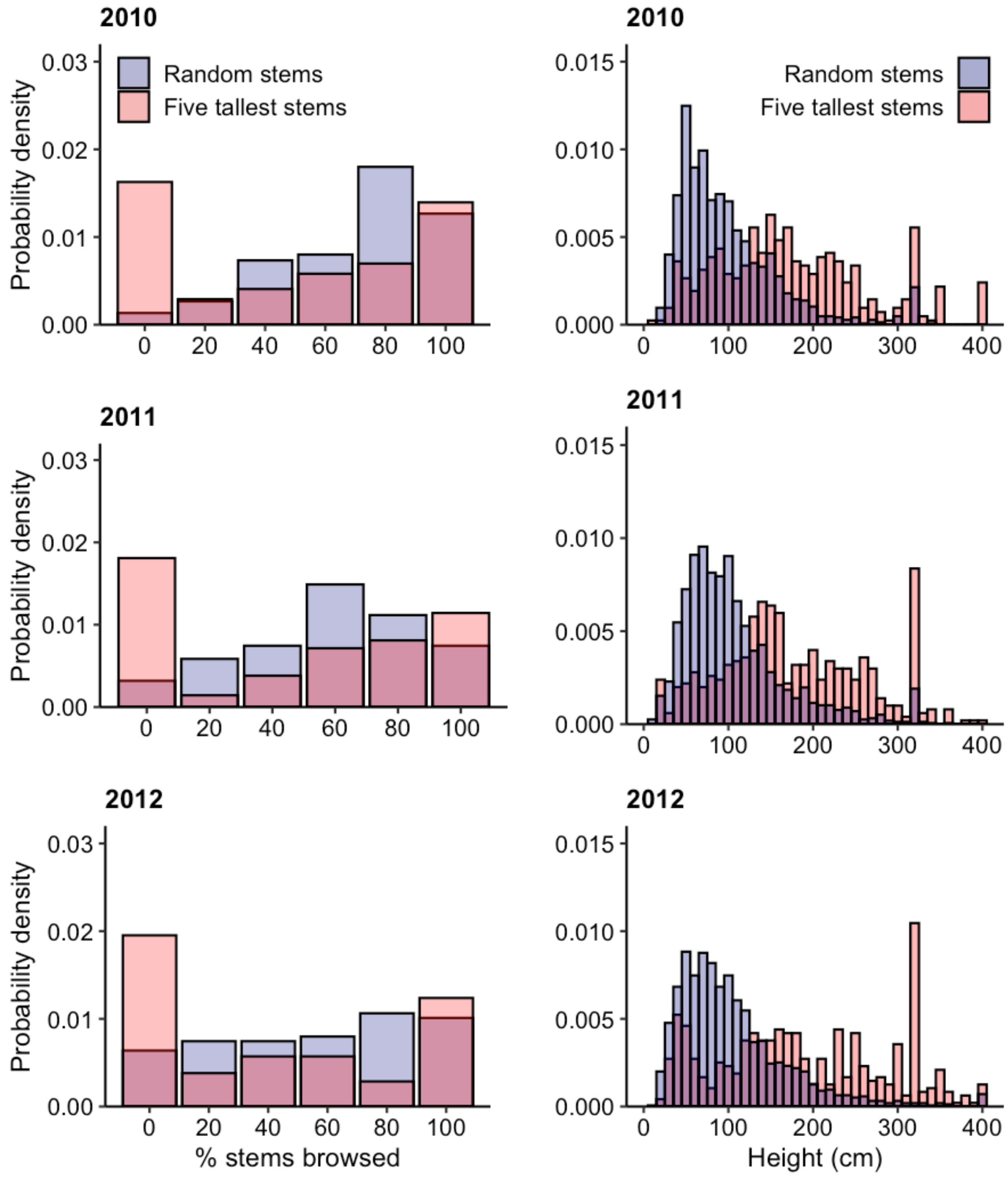

### Figure S2 continued

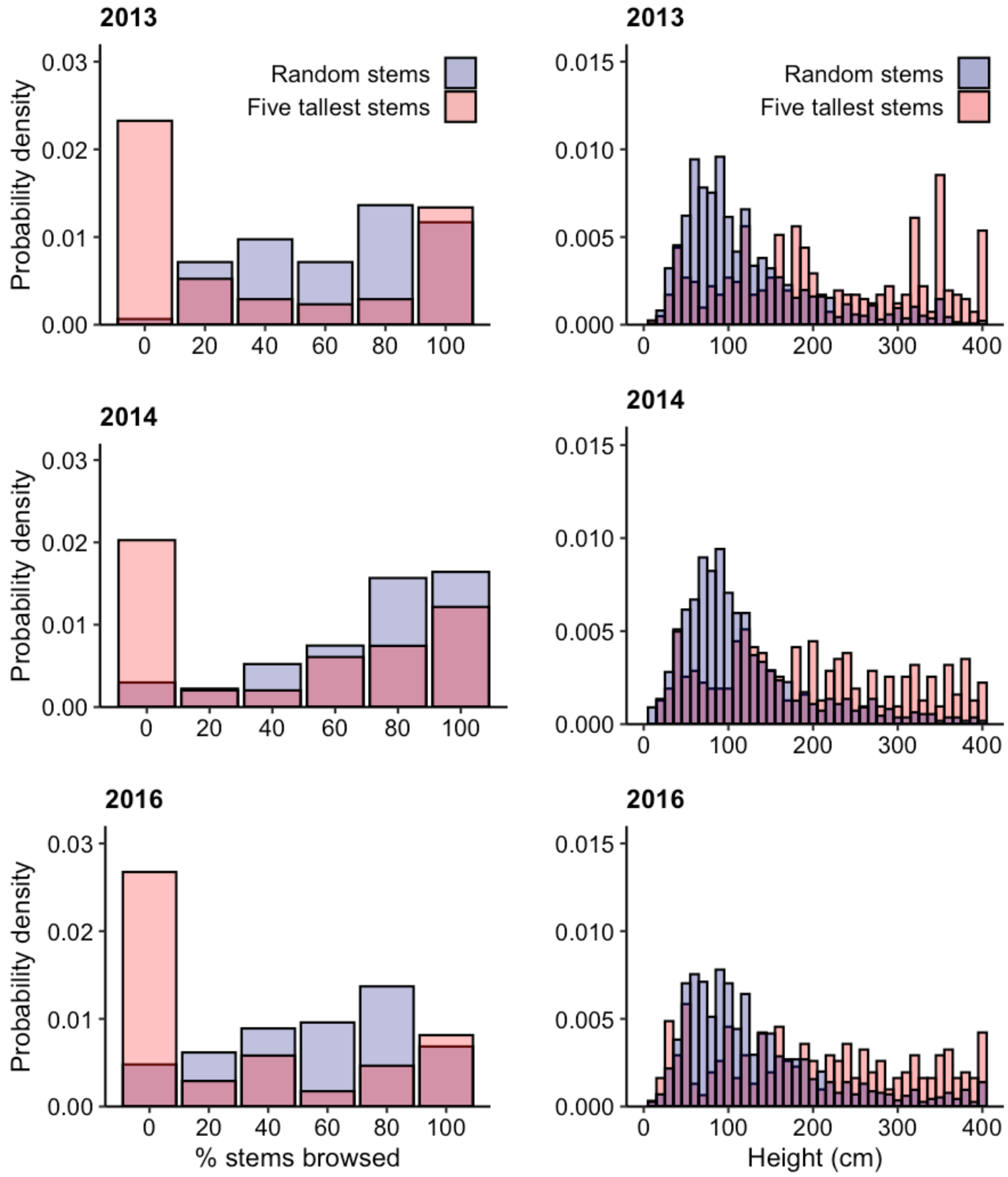

### Figure S2 continued

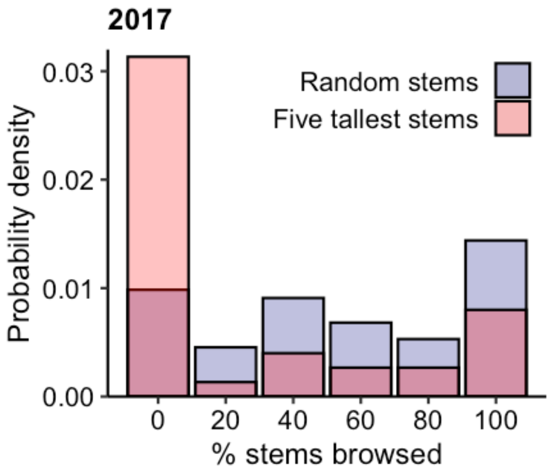


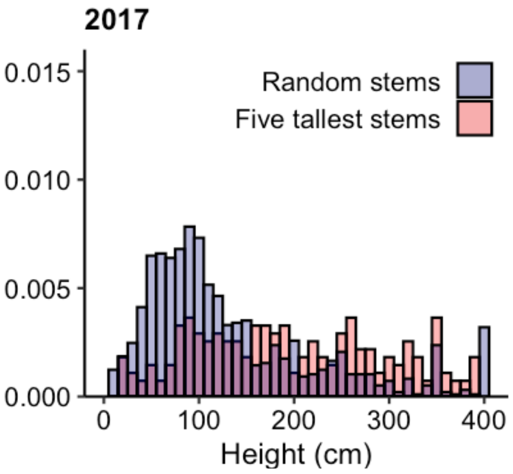

**Figure S3.** Annual trends in height percentiles of randomly selected young aspen (lines) and the five tallest young aspen (dots) pooled across 113 plots and stands. Median height ($50^{th}$ percentile) of the five tallest young aspen tracked the $85\text{-}90^{th}$ height percentile of random young aspen.

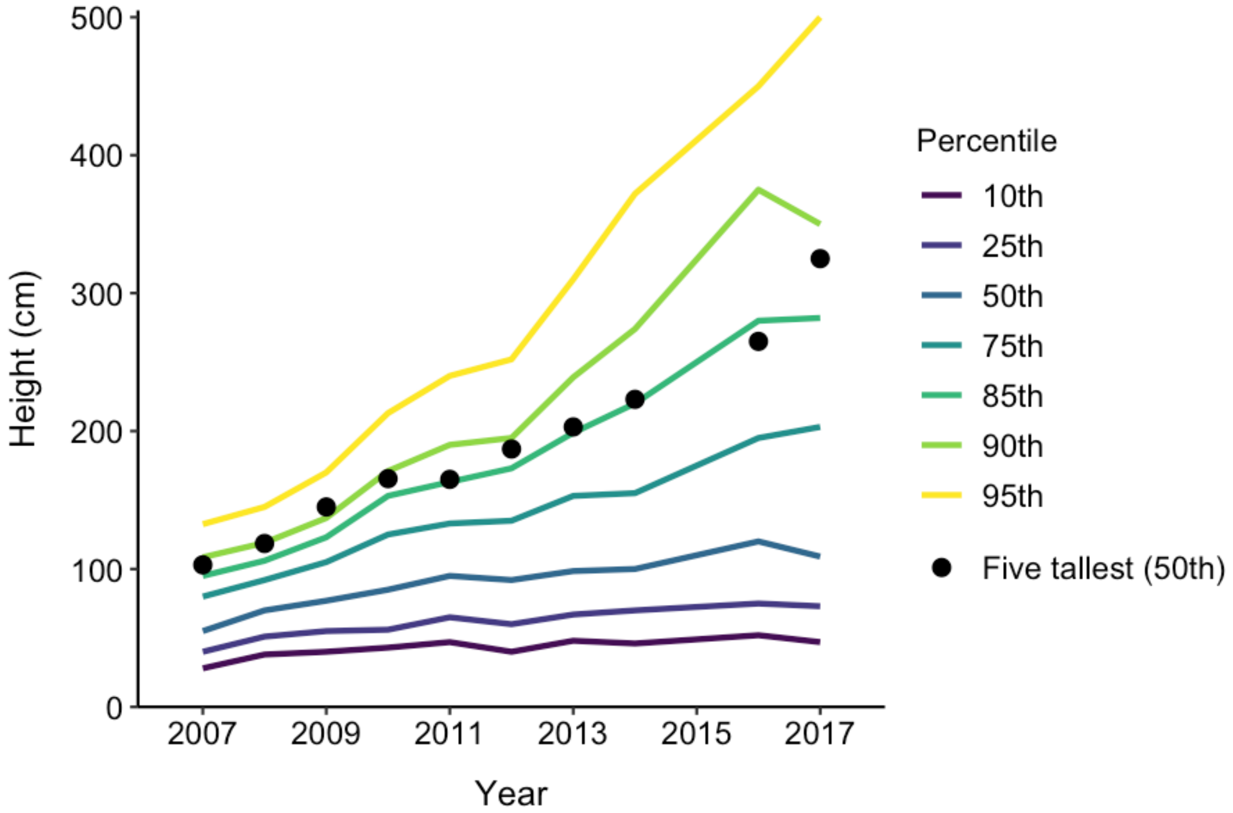

**Figure S4.** Annual percentage of randomly sampled aspen plots (a) and stands (b) with no young aspen. Eight plots had no young aspen for the entire study, but every stand had at least one year with young aspen. In 2007, 14 plots and 10 stands had no stems; of these, 10 plots (71.4%) and 2 stands (20%) also had no stems in 2017. Numbers above bars indicate the number of plots/stands measured each year.

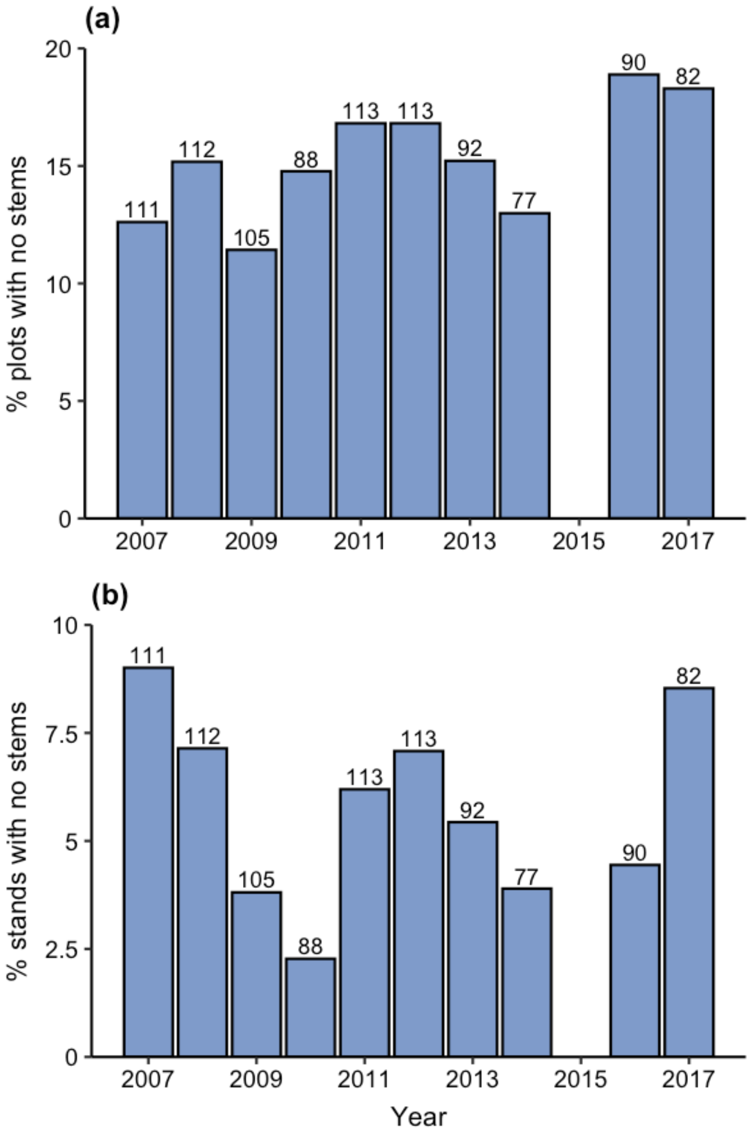

**Figure S5.** Photographs from 2001/2009 and 2017 of four sampled aspen stands with varying levels of young aspen regeneration. There was a persistent lack of regeneration in Plots 69, 81, and 85 between 2001/2009 (a, c, e) and 2017 (b, d, f). By contrast, Plot 72 exhibited a substantial increase in regeneration between 2009 (g) and 2017 (h). Plots 69 and 72 were within 1 km of each other.

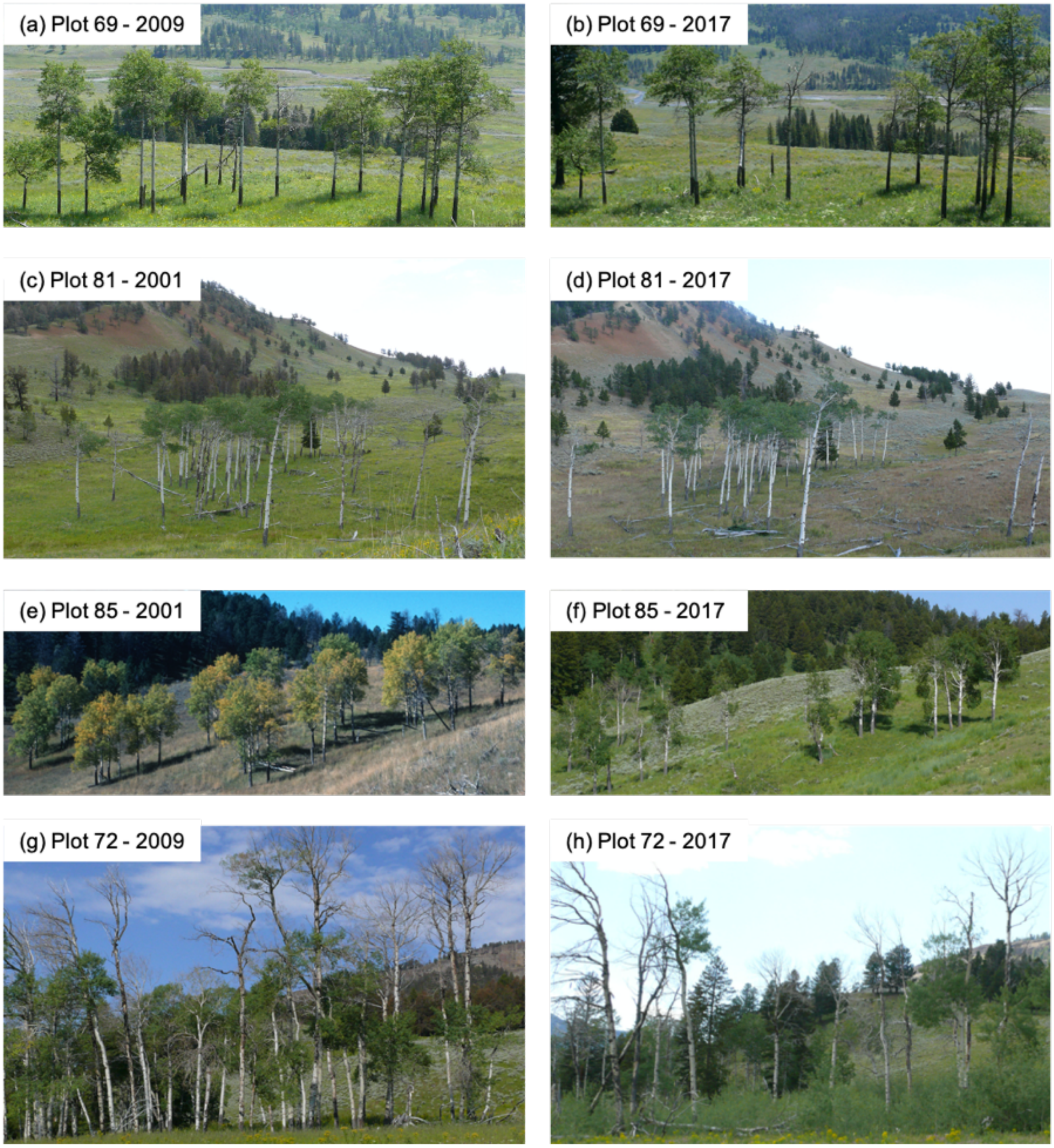

**Figure S6.** Elk in northern Yellowstone National Park prefer to browse young aspen at approximately the level of their shoulder height (~119-168 cm). Photo at left: Plot 58 (44.879828° N, 110.137308° W); December 19, 2019; 23:22 hours. Photo at right: Plot 14 (44.947878° N, 110.387356° W); November 8, 2019; 15:37 hours.

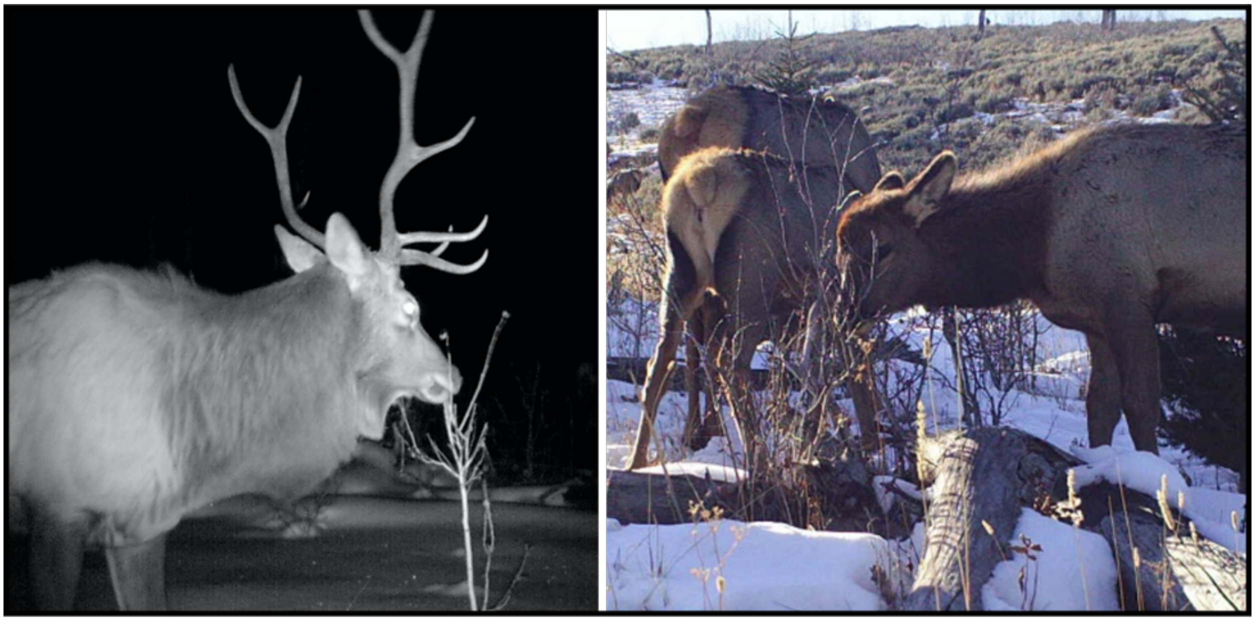

**Figure S7.** Example of the loss of a non-sampled aspen stand in northern Yellowstone National Park despite the reintroduction of wolves. This stand was located on the western shore of the Lamar River (44.853932° N, 110.192110° W). The 1954 and 1992 images are aerial photographs from Larsen and Ripple (2005) and the 2015 image is a satellite product from Google Earth (earth.google.com/web/). Dead and downed aspen trees are visible on the ground as white lines in the 1992 image and are also visible in the 2015 image when viewed at a larger scale in Google Earth. The area did not burn in the interval 1954-2015.

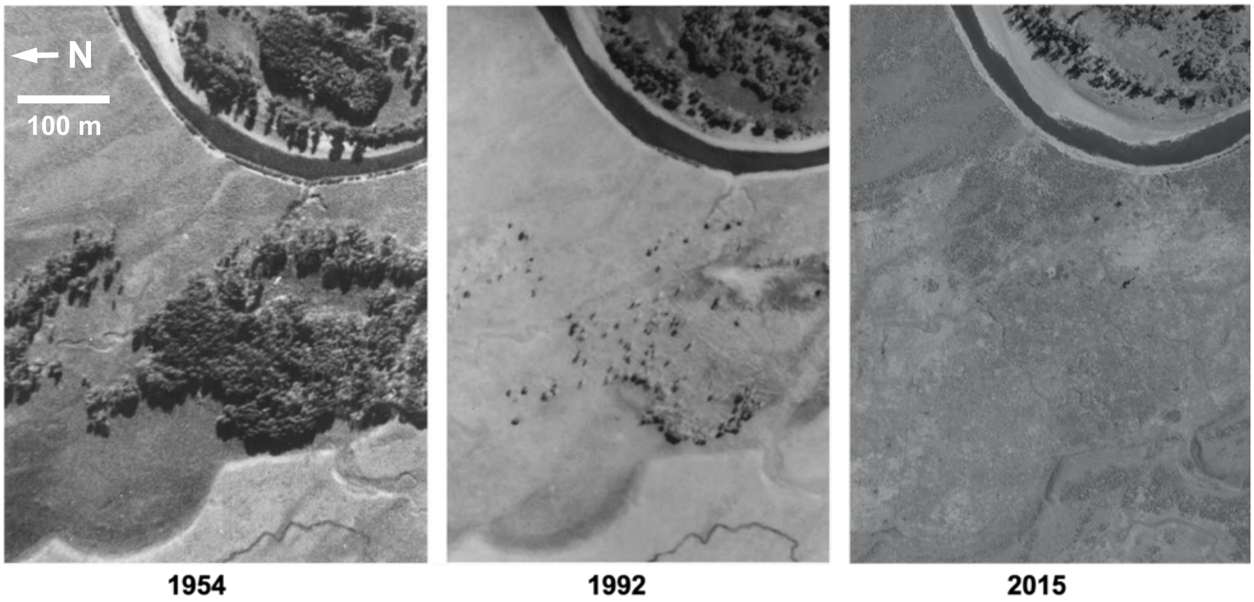

**Figure S8.** Distribution of plots with respect to projected changes in aspen habitat suitability by 2025, 2055, and 2085 (N.B. Piekielek, personal comm.; also see Piekielek *et al.* 2015 for projections to 2040, 2070, and 2099). Most of our sampled aspen stands occurred in or adjacent to areas projected to become unsuitable for aspen due to anthropogenic climate forcing consistent with increases in atmospheric greenhouse gases at rates similar to present. "No data" cells are areas not surveyed by the U.S. Forest Service Forest Inventory and Analysis program.

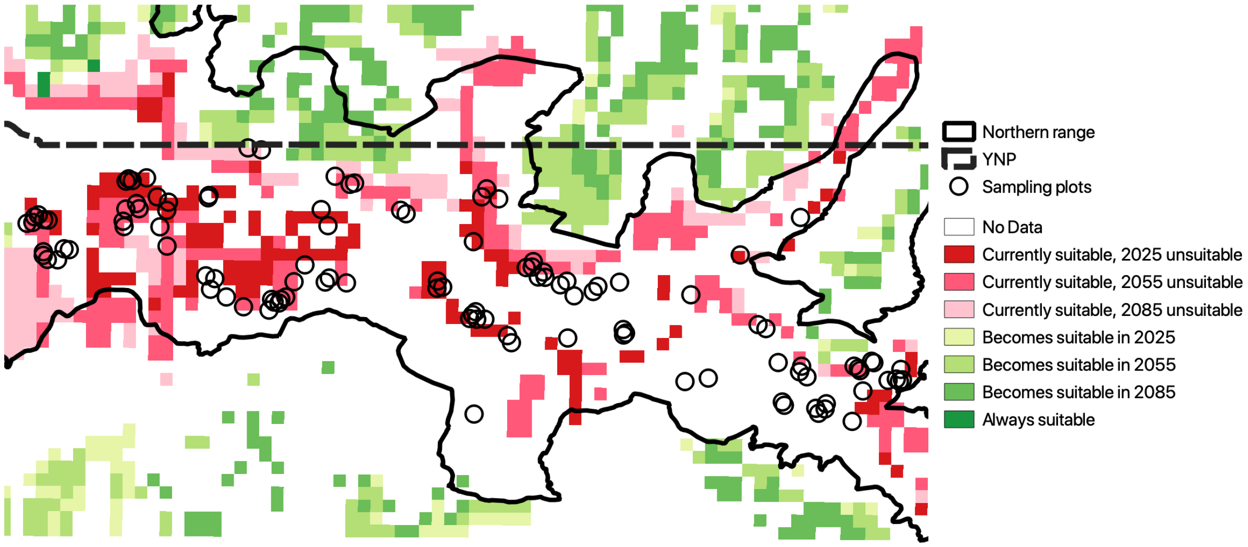

**Appendix S1**: Supplementary materials for "Sampling bias exaggerates a textbook example of a trophic cascade"

**Table S1.** Model selection results for GLMMs describing the effect of stem height (ht) on the probability that a "five tallest" young aspen was browsed in northern Yellowstone NP. Variables ht1 and ht2 contain a linear spline at the indicated knot (cm). The intercept and simple linear models included no knot. All models included crossed random effects for stand identity and year. Log-likelihood (LL), number of parameters (*K*), $AIC_c$, differences in $AIC_c$ compared to the best model ($\Delta AIC_c$), and $AIC_c$ weights (*W*) are given for each model. The best model ($\Delta AIC_c = 0.00$) is in boldface and competitive models ($\Delta AIC_c < 2.00$) are shaded.

| Model | Knot (cm) | LL | *K* | $AIC_c$ | $\Delta AIC_c$ | *W* |
|---|---|---|---|---|---|---|
| intercept | - | -2166.41 | 3 | 4339.04 | 749.59 | 0.00 |
| ht | - | -1845.00 | 4 | 3698.36 | 108.92 | 0.00 |
| ht1, ht2 | 10 | -1845.00 | 5 | 3700.55 | 111.11 | 0.00 |
| ht1, ht2 | 20 | -1839.17 | 5 | 3688.90 | 99.46 | 0.00 |
| ht1, ht2 | 30 | -1830.38 | 5 | 3671.32 | 81.87 | 0.00 |
| ht1, ht2 | 40 | -1822.30 | 5 | 3655.16 | 65.72 | 0.00 |
| ht1, ht2 | 50 | -1817.03 | 5 | 3644.62 | 55.17 | 0.00 |
| ht1, ht2 | 60 | -1813.72 | 5 | 3638.01 | 48.56 | 0.00 |
| ht1, ht2 | 70 | -1810.72 | 5 | 3632.01 | 42.57 | 0.00 |
| ht1, ht2 | 80 | -1808.52 | 5 | 3627.60 | 38.16 | 0.00 |
| ht1, ht2 | 90 | -1802.95 | 5 | 3616.47 | 27.03 | 0.00 |
| ht1, ht2 | 100 | -1797.70 | 5 | 3605.96 | 16.52 | 0.00 |
| ht1, ht2 | 110 | -1793.95 | 5 | 3598.46 | 9.01 | 0.00 |
| ht1, ht2 | 111 | -1793.71 | 5 | 3597.98 | 8.54 | 0.00 |
| ht1, ht2 | 112 | -1793.46 | 5 | 3597.48 | 8.04 | 0.00 |
| ht1, ht2 | 113 | -1793.19 | 5 | 3596.94 | 7.49 | 0.00 |
| ht1, ht2 | 114 | -1792.95 | 5 | 3596.47 | 7.03 | 0.00 |
| ht1, ht2 | 115 | -1792.75 | 5 | 3596.07 | 6.62 | 0.00 |
| ht1, ht2 | 116 | -1792.61 | 5 | 3595.78 | 6.34 | 0.00 |

Table S1 continued

| Model | Knot (cm) | LL | *K* | $AIC_c$ | $\Delta AIC_c$ | *W* |
|---|---|---|---|---|---|---|
| ht1, ht2 | 117 | -1792.47 | 5 | 3595.51 | 6.07 | 0.00 |
| ht1, ht2 | 118 | -1792.36 | 5 | 3595.27 | 5.83 | 0.00 |
| ht1, ht2 | 119 | -1792.17 | 5 | 3594.91 | 5.47 | 0.01 |
| ht1, ht2 | 120 | -1792.02 | 5 | 3594.59 | 5.15 | 0.01 |
| ht1, ht2 | 121 | -1791.77 | 5 | 3594.11 | 4.67 | 0.01 |
| ht1, ht2 | 122 | -1791.53 | 5 | 3593.62 | 4.18 | 0.01 |
| ht1, ht2 | 123 | -1791.28 | 5 | 3593.12 | 3.67 | 0.01 |
| ht1, ht2 | 124 | -1791.06 | 5 | 3592.68 | 3.24 | 0.02 |
| ht1, ht2 | 125 | -1790.89 | 5 | 3592.33 | 2.89 | 0.02 |
| ht1, ht2 | 126 | -1790.57 | 5 | 3591.69 | 2.25 | 0.03 |
| ht1, ht2 | 127 | -1790.27 | 5 | 3591.10 | 1.66 | 0.03 |
| ht1, ht2 | 128 | -1789.96 | 5 | 3590.49 | 1.04 | 0.05 |
| ht1, ht2 | 129 | -1789.70 | 5 | 3589.97 | 0.52 | 0.06 |
| ht1, ht2 | 130 | -1789.53 | 5 | 3589.63 | 0.18 | 0.07 |
| ht1, ht2 | 131 | -1789.47 | 5 | 3589.51 | 0.06 | 0.07 |
| **ht1, ht2** | **132** | **-1789.44** | **5** | **3589.44** | **0.00** | **0.08** |
| ht1, ht2 | 133 | -1789.48 | 5 | 3589.51 | 0.07 | 0.07 |
| ht1, ht2 | 134 | -1789.52 | 5 | 3589.61 | 0.16 | 0.07 |
| ht1, ht2 | 135 | -1789.61 | 5 | 3589.77 | 0.33 | 0.07 |
| ht1, ht2 | 136 | -1789.65 | 5 | 3589.86 | 0.41 | 0.06 |
| ht1, ht2 | 137 | -1789.75 | 5 | 3590.06 | 0.61 | 0.06 |
| ht1, ht2 | 138 | -1789.89 | 5 | 3590.35 | 0.90 | 0.05 |
| ht1, ht2 | 139 | -1790.07 | 5 | 3590.71 | 1.26 | 0.04 |
| ht1, ht2 | 140 | -1790.33 | 5 | 3591.23 | 1.79 | 0.03 |
| ht1, ht2 | 141 | -1790.66 | 5 | 3591.88 | 2.44 | 0.02 |
| ht1, ht2 | 142 | -1791.00 | 5 | 3592.56 | 3.11 | 0.02 |
| ht1, ht2 | 143 | -1791.37 | 5 | 3593.31 | 3.86 | 0.01 |

Table S1 continued

| Model | Knot (cm) | LL | $K$ | $AIC_c$ | $\Delta AIC_c$ | $W$ |
|---|---|---|---|---|---|---|
| ht1, ht2 | 144 | -1791.85 | 5 | 3594.25 | 4.81 | 0.01 |
| ht1, ht2 | 145 | -1792.34 | 5 | 3595.23 | 5.79 | 0.00 |
| ht1, ht2 | 146 | -1792.80 | 5 | 3596.15 | 6.71 | 0.00 |
| ht1, ht2 | 147 | -1793.25 | 5 | 3597.06 | 7.62 | 0.00 |
| ht1, ht2 | 148 | -1793.70 | 5 | 3597.97 | 8.53 | 0.00 |
| ht1, ht2 | 149 | -1794.21 | 5 | 3598.98 | 9.54 | 0.00 |
| ht1, ht2 | 150 | -1794.70 | 5 | 3599.97 | 10.53 | 0.00 |
| ht1, ht2 | 160 | -1799.25 | 5 | 3609.06 | 19.62 | 0.00 |
| ht1, ht2 | 170 | -1803.13 | 5 | 3616.83 | 27.39 | 0.00 |
| ht1, ht2 | 180 | -1807.42 | 5 | 3625.39 | 35.95 | 0.00 |
| ht1, ht2 | 190 | -1811.10 | 5 | 3632.77 | 43.33 | 0.00 |
| ht1, ht2 | 200 | -1813.79 | 5 | 3638.13 | 48.69 | 0.00 |

**Table S2.** Model selection results for GLMMs describing the effect of stem height (ht) on the probability that a randomly sampled young aspen was browsed in northern Yellowstone NP. Variables ht1 and ht2 contain a linear spline at the indicated knot (cm). The intercept and simple linear models included no knot. All models included crossed random effects for stand identity and year. Log-likelihood (LL), number of parameters (*K*), $AIC_c$, differences in $AIC_c$ compared to the best model ($\Delta AIC_c$), and $AIC_c$ weights (*W*) are given for each model. The best model ($\Delta AIC_c$ = 0.00) is in boldface and competitive models ($\Delta AIC_c$ < 2.00) are shaded.

| Model | Knot (cm) | LL | *K* | $AIC_c$ | $\Delta AIC_c$ | *W* |
|---|---|---|---|---|---|---|
| intercept | - | -8238.90 | 3 | 16484.05 | 1028.55 | 0.00 |
| ht | - | -7886.08 | 4 | 15780.57 | 325.08 | 0.00 |
| ht1, ht2 | 10 | -7885.74 | 5 | 15782.08 | 326.59 | 0.00 |
| ht1, ht2 | 20 | -7886.00 | 5 | 15782.60 | 327.11 | 0.00 |
| ht1, ht2 | 30 | -7883.24 | 5 | 15777.08 | 321.58 | 0.00 |
| ht1, ht2 | 40 | -7877.04 | 5 | 15764.69 | 309.20 | 0.00 |
| ht1, ht2 | 50 | -7859.22 | 5 | 15729.05 | 273.56 | 0.00 |
| ht1, ht2 | 60 | -7835.44 | 5 | 15681.48 | 225.99 | 0.00 |
| ht1, ht2 | 70 | -7803.77 | 5 | 15618.14 | 162.65 | 0.00 |
| ht1, ht2 | 80 | -7776.04 | 5 | 15562.69 | 107.19 | 0.00 |
| ht1, ht2 | 90 | -7755.96 | 5 | 15522.53 | 67.04 | 0.00 |
| ht1, ht2 | 100 | -7738.23 | 5 | 15487.07 | 31.58 | 0.00 |
| ht1, ht2 | 110 | -7726.73 | 5 | 15464.07 | 8.58 | 0.00 |
| ht1, ht2 | 111 | -7726.10 | 5 | 15462.81 | 7.32 | 0.00 |
| ht1, ht2 | 112 | -7725.49 | 5 | 15461.59 | 6.10 | 0.00 |
| ht1, ht2 | 113 | -7724.91 | 5 | 15460.43 | 4.94 | 0.01 |
| ht1, ht2 | 114 | -7724.41 | 5 | 15459.43 | 3.94 | 0.01 |
| ht1, ht2 | 115 | -7724.08 | 5 | 15458.78 | 3.28 | 0.02 |
| ht1, ht2 | 116 | -7723.73 | 5 | 15458.07 | 2.58 | 0.03 |

Table S2 continued

| Model | Knot (cm) | LL | *K* | $AIC_c$ | $\Delta AIC_c$ | *W* |
|---|---|---|---|---|---|---|
| ht1, ht2 | 117 | -7723.49 | 5 | 15457.58 | 2.08 | 0.03 |
| ht1, ht2 | 118 | -7723.15 | 5 | 15456.91 | 1.42 | 0.05 |
| ht1, ht2 | 119 | -7722.91 | 5 | 15456.43 | 0.94 | 0.06 |
| ht1, ht2 | 120 | -7722.62 | 5 | 15455.85 | 0.35 | 0.08 |
| ht1, ht2 | 121 | -7722.51 | 5 | 15455.63 | 0.14 | 0.09 |
| **ht1, ht2** | **122** | **-7722.44** | **5** | **15455.49** | **0.00** | **0.09** |
| ht1, ht2 | 123 | -7722.47 | 5 | 15455.54 | 0.05 | 0.09 |
| ht1, ht2 | 124 | -7722.55 | 5 | 15455.70 | 0.20 | 0.08 |
| ht1, ht2 | 125 | -7722.68 | 5 | 15455.97 | 0.47 | 0.07 |
| ht1, ht2 | 126 | -7722.81 | 5 | 15456.22 | 0.73 | 0.06 |
| ht1, ht2 | 127 | -7722.99 | 5 | 15456.58 | 1.09 | 0.05 |
| ht1, ht2 | 128 | -7723.19 | 5 | 15456.98 | 1.49 | 0.04 |
| ht1, ht2 | 129 | -7723.39 | 5 | 15457.39 | 1.90 | 0.04 |
| ht1, ht2 | 130 | -7723.66 | 5 | 15457.92 | 2.42 | 0.03 |
| ht1, ht2 | 131 | -7723.99 | 5 | 15458.60 | 3.10 | 0.02 |
| ht1, ht2 | 132 | -7724.36 | 5 | 15459.33 | 3.84 | 0.01 |
| ht1, ht2 | 133 | -7724.81 | 5 | 15460.24 | 4.74 | 0.01 |
| ht1, ht2 | 134 | -7725.26 | 5 | 15461.12 | 5.63 | 0.01 |
| ht1, ht2 | 135 | -7725.81 | 5 | 15462.22 | 6.72 | 0.00 |
| ht1, ht2 | 136 | -7726.37 | 5 | 15463.35 | 7.86 | 0.00 |
| ht1, ht2 | 137 | -7727.03 | 5 | 15464.66 | 9.17 | 0.00 |
| ht1, ht2 | 138 | -7727.75 | 5 | 15466.11 | 10.62 | 0.00 |
| ht1, ht2 | 139 | -7728.53 | 5 | 15467.68 | 12.18 | 0.00 |
| ht1, ht2 | 140 | -7729.36 | 5 | 15469.33 | 13.84 | 0.00 |
| ht1, ht2 | 141 | -7730.18 | 5 | 15470.96 | 15.47 | 0.00 |
| ht1, ht2 | 142 | -7731.08 | 5 | 15472.76 | 17.27 | 0.00 |
| ht1, ht2 | 143 | -7732.01 | 5 | 15474.64 | 19.14 | 0.00 |

Table S2 continued

| Model | Knot (cm) | LL | *K* | $AIC_c$ | $\Delta AIC_c$ | *W* |
|---|---|---|---|---|---|---|
| ht1, ht2 | 144 | -7733.11 | 5 | 15476.82 | 21.33 | 0.00 |
| ht1, ht2 | 145 | -7734.31 | 5 | 15479.23 | 23.74 | 0.00 |
| ht1, ht2 | 146 | -7735.54 | 5 | 15481.68 | 26.19 | 0.00 |
| ht1, ht2 | 147 | -7736.72 | 5 | 15484.05 | 28.56 | 0.00 |
| ht1, ht2 | 148 | -7737.86 | 5 | 15486.32 | 30.83 | 0.00 |
| ht1, ht2 | 149 | -7739.02 | 5 | 15488.64 | 33.15 | 0.00 |
| ht1, ht2 | 150 | -7740.22 | 5 | 15491.05 | 35.56 | 0.00 |
| ht1, ht2 | 160 | -7749.68 | 5 | 15509.98 | 54.48 | 0.00 |
| ht1, ht2 | 170 | -7760.17 | 5 | 15530.95 | 75.46 | 0.00 |
| ht1, ht2 | 180 | -7771.68 | 5 | 15553.96 | 98.47 | 0.00 |
| ht1, ht2 | 190 | -7781.88 | 5 | 15574.36 | 118.86 | 0.00 |
| ht1, ht2 | 200 | -7793.35 | 5 | 15597.31 | 141.82 | 0.00 |
